# Deletion of the endothelial glycocalyx component endomucin leads to impaired glomerular structure and function

**DOI:** 10.1101/2024.07.16.603749

**Authors:** Zhengping Hu, Issahy Cano, Fengyang Lei, Jie Liu, Ramon Bossardi Ramos, Harper Gordon, Eleftherios I. Paschalis, Magali Saint-Geniez, Yin Shan Eric Ng, Patricia A. D’Amore

**Affiliations:** Schepens Eye Research Institute of Massachusetts Eye and Ear, Boston, Massachusetts; Departments of Ophthalmology and Pathology, Harvard Medical School, Boston, Massachusetts; Departments of Pathology, Harvard Medical School, Boston, Massachusetts; Department of Molecular and Cellular Physiology, Albany Medical Center, Albany, New York

**Keywords:** glycocalyx, vascular endothelial growth factor (VEGF), inflammation, podocyte, glomerular filtration barrier

## Abstract

**Background:** Endomucin (EMCN), an endothelial-specific glycocalyx component, was found to be highly expressed by the endothelium of the renal glomerulus. We reported an anti-inflammatory role of EMCN and its involvement in the regulation of vascular endothelial growth factor (VEGF) activity through modulating VEGF receptor 2 (VEGFR2) endocytosis. The goal of this study is to investigate the phenotypic and functional effects of EMCN deficiency using the first global EMCN knockout mouse model.

**Methods:** Global EMCN knockout mice were generated by crossing EMCN-floxed mice with ROSA26-Cre mice. Flow cytometry was employed to analyze infiltrating myeloid cells in the kidneys. The ultrastructure of the glomerular filtration barrier was examined by transmission electron microscopy, while urinary albumin, creatinine, and total protein levels were analyzed from freshly collected urine samples. Expression and localization of EMCN, EGFP, CD45, CD31, CD34, podocin, albumin, and α-smooth muscle actin were examined by immunohistochemistry. Mice were weighed regularly, and their systemic blood pressure was measured using a non-invasive tail-cuff system. Glomerular endothelial cells and podocytes were isolated by fluorescence-activated cell sorting for RNA-seq. Transcriptional profiles were analyzed to identify differentially expressed genes in both endothelium and podocytes, followed by gene ontology analysis of up- and down-regulated genes. Protein levels of EMCN, albumin, and podocin were quantified by Western blot.

**Results:** EMCN^-/-^ mice were viable with no gross anatomical defects in kidneys. The EMCN^-/-^ mice exhibited increased infiltration of CD45^+^ cells, with an increased proportion of Ly6G^high^Ly6C^high^ myeloid cells and higher VCAM-1 expression. EMCN^-/-^ mice displayed albuminuria with increased albumin in the Bowman’s space compared to the EMCN^+/+^ littermates. Glomeruli in EMCN^-/-^ mice revealed fused and effaced podocyte foot processes and disorganized endothelial fenestrations. We found no significant difference in blood pressure between EMCN knockout mice and their wild-type littermates. RNA-seq of glomerular endothelial cells revealed downregulation of cell-cell adhesion and MAPK/ERK pathways, along with glycocalyx and extracellular matrix remodeling. In podocytes, we observed reduced VEGF signaling and alterations in cytoskeletal organization. Notably, there was a significant decrease in both mRNA and protein levels of podocin, a key component of the slit diaphragm.

**Conclusion:** Our study demonstrates a critical role of the endothelial marker EMCN in supporting normal glomerular filtration barrier structure and function by maintaining glomerular endothelial tight junction and homeostasis and podocyte function through endothelial-podocyte crosstalk.

## INTRODUCTION

The endothelial glycocalyx is a structure that emerges from the luminal surface of the vascular endothelium, which lines the entire vasculature. It is a negatively charged, carbohydrate-rich structure composed of proteoglycans, glycoproteins, glycolipids, and glycosaminoglycans, such as heparan sulfate, chondroitin sulfate, hyaluronan, along with associated plasma proteins. Due to its structural complexity, the functional role of endothelial glycocalyx has been largely unexplored. Observations made over the past several decades have pointed to a critical role for the glycocalyx in modulating vascular inflammation, permeability, signaling, endothelial homeostasis, and in maintaining the integrity of the endothelial barrier^1-4^. Disruption of the glycocalyx is observed in a variety of pathologies, including sepsis, atherosclerosis^5^, kidney disease^6-8^, and diabetes^9,10^. Endomucin (EMCN), a component of the endothelial glycocalyx, was first identified in 1999 as an endothelial-specific sialomucin^11^. It is a type I transmembrane glycoprotein that is heavily O-glycosylated on its extracellular domain and selectively expressed by venous as well as capillary endothelium but not by arterial endothelium^11,12^. Increasing evidence has revealed important functional roles for EMCN in hematopoietic stem cell differentiation, inflammation, and angiogenesis^12-15^. Overexpression of EMCN has also been reported to promote cell detachment and to interfere with the assembly of the focal adhesion complex, indicating a role of EMCN in endothelial anti-adhesive activity^16^. EMCN isolated from human and mouse lymphoid organs was found decorated with a specific epitope, MECA-79, which defines L-selectin ligands^9,17^ suggesting that EMCN can function as a ligand for L-selectin^9-18^ to mediate and support L-selectin-dependent leukocyte rolling under flow conditions. Prior investigations of the role of EMCN in leukocyte-endothelial interactions by our group indicated that knockdown of EMCN in quiescent endothelial cells (ECs) leads to robust adhesion of neutrophils. This adhesion occurs through LFA-1-mediated interaction with intercellular adhesion molecule 1 (ICAM-1), which is consistently expressed in vitro by ECs^14^. Moreover, overexpression of EMCN significantly suppressed leukocyte adhesion induced by tumor necrosis factor-alpha (TNF-α) in a mouse model of ocular inflammation, suggesting a role of EMCN as a crucial modulator of leukocyte infiltration^14^.

EMCN has also been shown to label hematopoietic stem cells throughout development^13^ and has been suggested to play a role in angiogenesis as that upregulation of EMCN occurs in ECs stimulated to proliferate with tumor-conditioned media^19^. We have shown that knockdown of EMCN in vivo leads to impaired angiogenesis during retinal development and significantly inhibits vascular endothelial growth factor (VEGF)-induced cell migration, tube formation, and proliferation in vitro^15^. This effect is mediated by EMCN’s role in regulating clathrin-mediated endocytosis of activated VEGF receptor 2 (VEGFR2) and downstream signaling upon VEGF stimulation through its glycosylated extracellular domain^12,20^.

To further define the functional role of EMCN in endothelial biology, global EMCN knockout (KO) mice were generated. Based on the relatively high levels of EMCN expression detected in the vasculature of the kidney, we prioritized and focused our initial investigation on this new KO model by investigating the structural and functional changes of EMCN deletion in the kidney. We observed that the kidneys of EMCN KO mice displayed a variety of defects, including increased inflammatory cell infiltration, albuminuria, thickened glomerular basement membrane, as well as dysregulated capillary fenestrations, and podocyte effacement. These findings highlight a critical role for EMCN in the maintenance of kidney vascular permeability and the glomerular filtration barrier to support normal kidney function.

## METHODS

### Animals

Experimental animal procedures were performed in accordance with the guidelines and under the approval of the Institutional Animal Care and Use Committee of the Schepens Eye Research Institute of Massachusetts Eye and Ear. All animals received food and water ad libitum, with a 12-hour day-night cycle and in a temperature-controlled environment. EMCN-floxed mice were generated in collaboration with Cyagen Biosciences, using the strategy summarized in Figure 1A. In brief, loxP sites were inserted into the EMCN gene to flank the exons 2 to 10. A promoter-less reporter EGFP (in reverse orientation) was flanked by lox2272 sites and loxP sites and was expressed under the control of EMCN promoter following successful inversion by Cre recombination and deletion of exons 2-10 of the EMCN gene. Global homozygous EMCN KO (EMCN^-/-^) mice were obtained by crossing heterozygous EMCN^+/-^ mice that were generated by crossing EMCN^flox/flox^ mice with the ROSA26-Cre mice, as illustrated in Supplemental Figure 1A. Genomic DNA and RNA were extracted from EMCN^+/-^ and EMCN^-/-^ mice (8 to 16-week-old) to validate the deletion of EMCN and RNA reduction in various tissues, including ear snip, kidney, lung, thyroid, spleen, liver, and small intestine. The body weights of the mice were monitored every week up to 16 weeks and every other week from 16 to 24 weeks of age. Kidneys from 12 to 24-week-old mice were collected for immunohistochemistry, morphologic examinations, transmission electron microscopy (TEM), protein and RNA analysis, and kidney function analysis. All studies were conducted on littermates.

**Figure 1.**
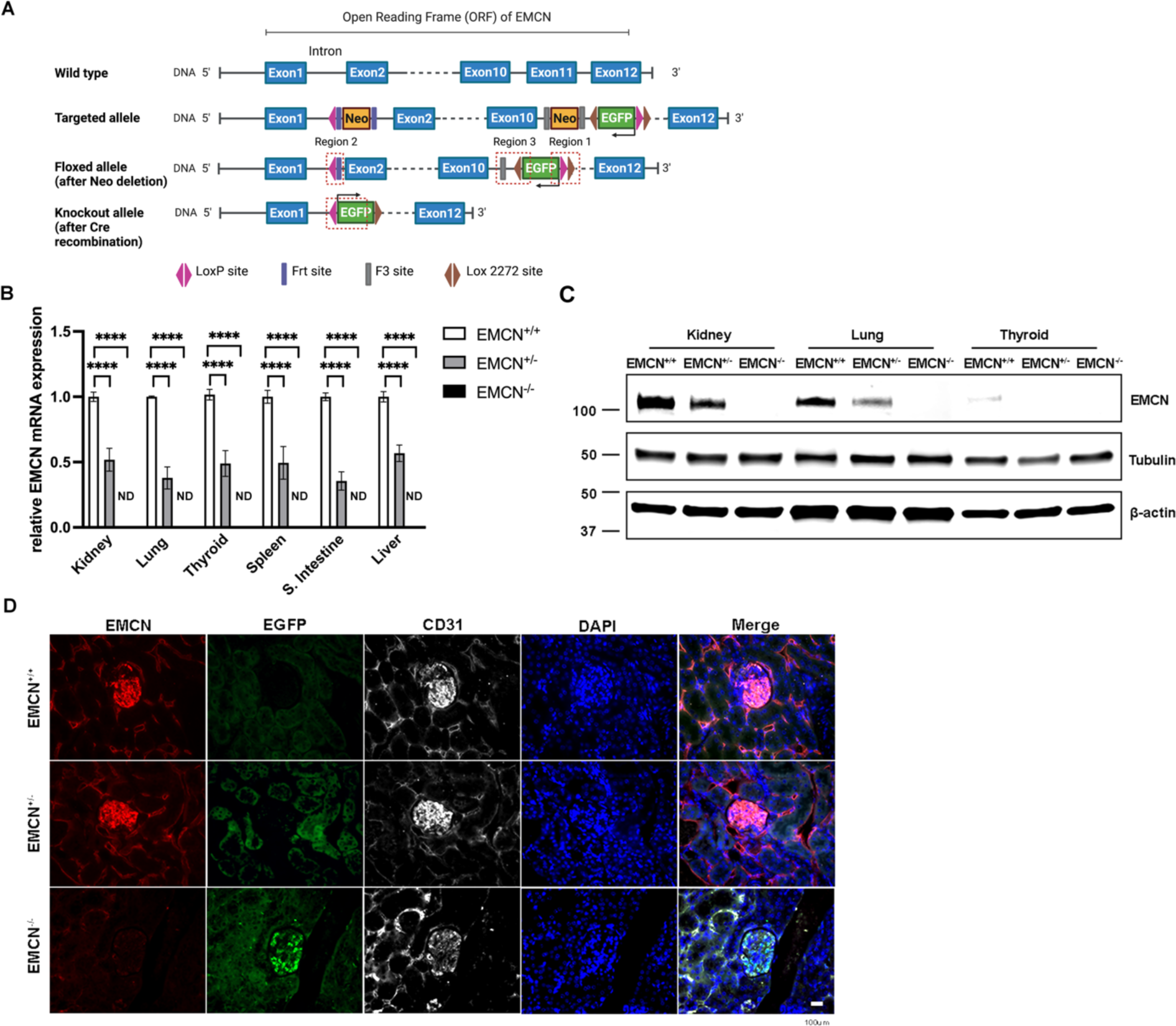
Generation of EMCN global knockout mice and confirmation of EMCN deletion. **(A)** EMCN floxed allele and EMCN deletion allele were generated as shown. The open reading frame (ORF) of EMCN was targeted and floxed by loxP and lox2272 sites with an EGFP tag. After Cre recombination, the majority region of EMCN ORF from exon 2 to exon 11 was deleted and the EGFP sequence was flipped, and resulting expression reflected EMCN deletion. **(B)** Kidney, lung, thyroid gland, spleen, small intestine, and liver tissues were collected from EMCN^+/+^, EMCN^+/-^, and EMCN^-/-^ mice (8 to 16 weeks old) and mRNA was extracted and examined for EMCN expression. ND indicated EMCN mRNA was not detected in EMCN^-/-^ tissues. n ≥ 4, **** *p* < 0.0001, one-way ANOVA. **(C)** Total protein was extracted from kidney, lung, and thyroid gland from EMCN^+/+^, EMCN^+/-^, and EMCN^-/-^ mice. Protein concentration was quantified using the BCA protein assay kit and 30 μg of each sample was loaded onto an SDS-PAGE gel. Levels of EMCN, tubulin, and β-actin were examined by western blot. n=3. **(D)** Kidneys from EMCN^+/+^, EMCN^+/-^, and EMCN^-/-^ mice were collected and fixed with 10% formalin overnight for paraffin sectioning. Immunohistochemical staining was performed to visualize EMCN, EGFP, and CD31; nuclei were stained with DAPI. n ≥ 4. Scale bar = 100 μm.

### Genotyping and sequence analysis

Genomic DNA was extracted from ear snips using QIAamp® Fast DNA Tissue Kit (Qiagen # 51404). Region 2 for wild-type or floxed EMCN gene, as well as Region 3 for KO of EMCN gene, as illustrated in Figure 1A, was amplified through polymerase chain reaction using specific primers (5′-AGTGAGTAGAGCATGGATTTGGGA-3′ wild-type or floxed allele-specific, 5′-CGATCACATGGTCCTGCTGGAGTT-3′ KO allele-specific, and 5′-AACATCCACTCTCTAGAGACCACACAT-3′ common primer). The wild-type allele resolves as a 259-bp band, the floxed allele resolves as a 424-bp band, and the KO allele resolves as a 663-bp band with agarose gel electrophoresis as the representative figure indicates (Supplemental Figure 1B). The 663-bp EMCN KO band was excised from agarose gel and sequenced, confirming the expression of the EGFP gene and deletion of the EMCN gene (Supplemental Figure 1C).

### RNA extraction, RT-PCR, and qPCR

Kidneys were collected in RNAlater (Invitrogen, #AM7020), and RNA was extracted using the RNeasy Mini Kit (Qiagen, #74136). Five hundred ng of purified RNA was reverse transcribed into cDNA using the SuperScript IV reverse transcriptase kit (Invitrogen, #11756050). Using FastStart Universal SYBR Green master mix (Roche, #A25743), gene expression was read with the Precision Plus Protein^TM^ LightCycler^®^ 480 II (Roche, Indianapolis, IN, United States) on 384 well plates containing a starting amount of 10-20 ng of cDNA. The primers used for RT-PCR are listed in Table 1. Relative mRNA level is quantified as Delta Delta Ct by first normalized to the average of 3 house-keeping genes: Hprt1, B2m, and Gapdh, and then normalized the resulting Delta Ct values to that from the wildtype control. All samples were prepared with the same amount of total RNA and volume of starting reagents for qPCR analysis.

**Table 1.**
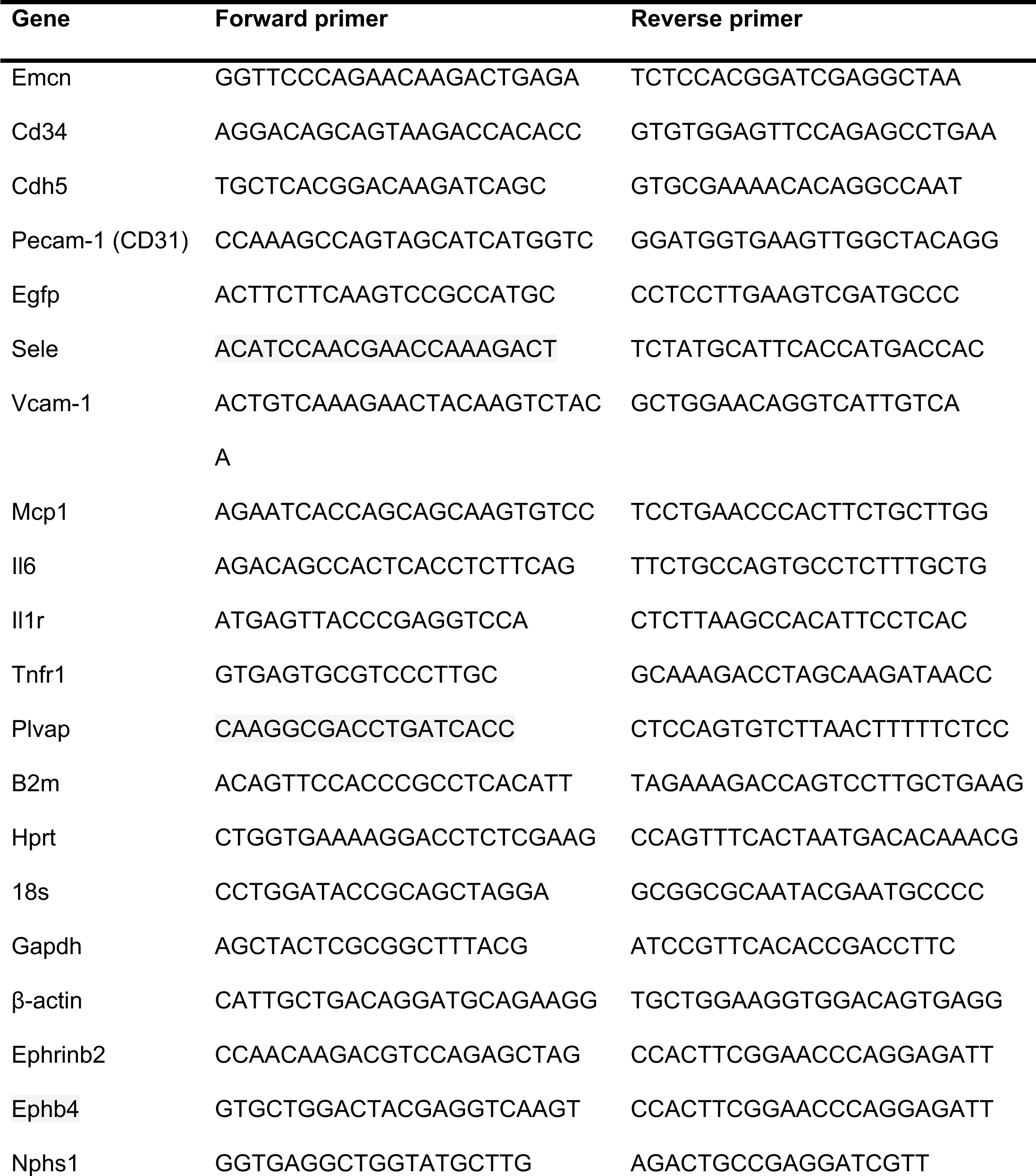
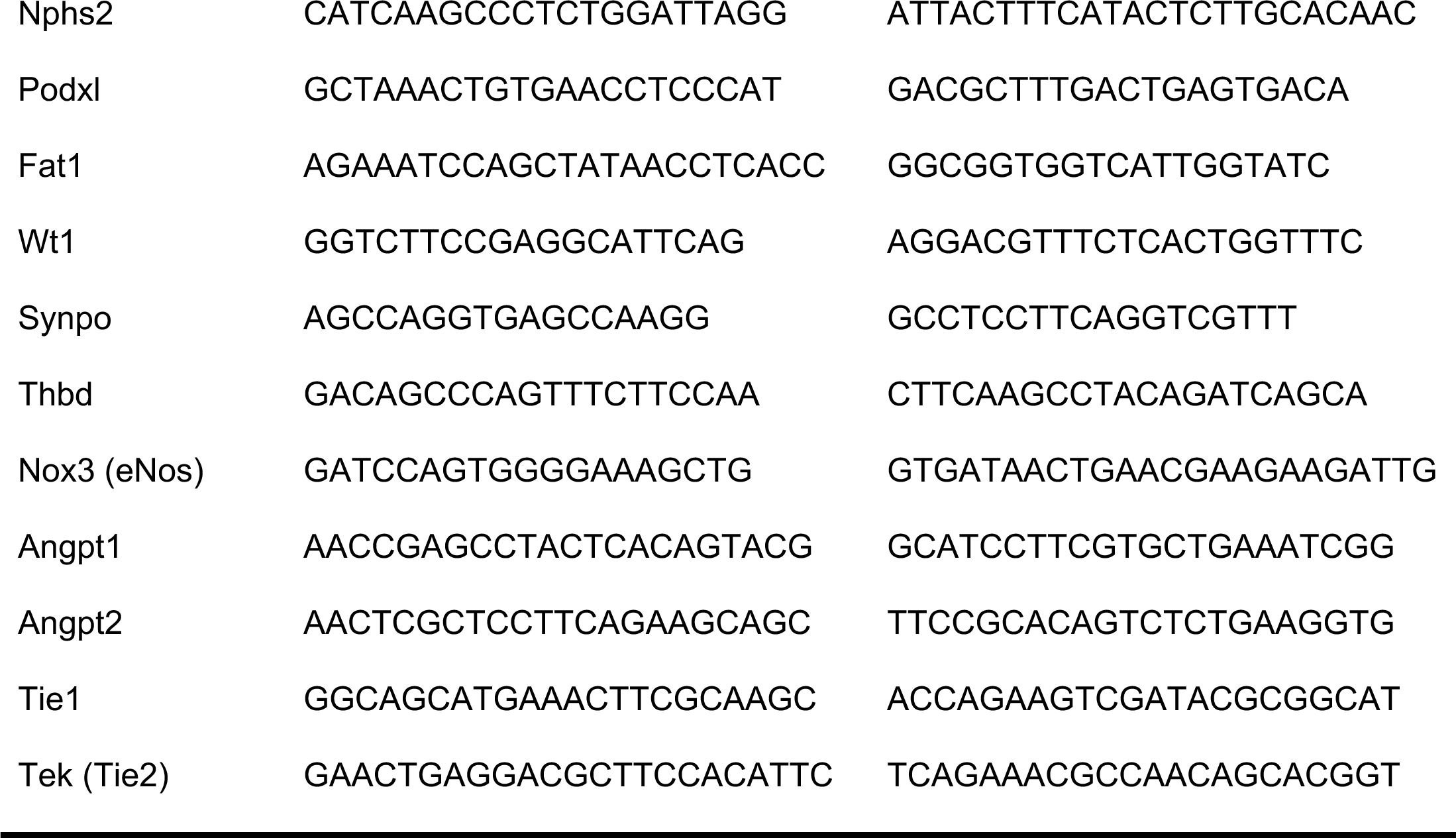
Sequence of primers used for RT-PCR studies.

### Reagents and antibodies

Lysis Buffer (#9803S) and protease inhibitors (#5871S) were purchased from Cell Signaling. Tween-20 (#X251-07), phosphate-buffered saline (PBS, #D5652-10x1L), DL-Dithiothreitol (DTT, #D9163-5G), and bovine serum albumin (BSA, #A6003) were purchased from Sigma-Aldrich. Precision Plus Protein Dual Color Standards (#161-0374) was obtained from Bio-Rad. BioTrace™ NT Nitrocellulose Transfer Membrane 30 cm x 3 m roll (#27376-991), DNA Gel Loading Dye (#R0611), and Fisherbrand™ RNase-Free Disposable Pellet Pestles (#12-141-364) were purchased from Fisher Scientific. Invitrogen™ RNAlater™ Stabilization Solution (#AM7020) was purchased from Invitrogen. RNase-Free DNase Set (#79254), RNeasy plus mini kit (#74136), and QIAamp Fast DNA Tissue Kit (#51404) were purchased from Qiagen.

Immunoblots were probed with rat anti-mouse EMCN (1:1,000, Abcam, #ab106100), mouse anti-tubulin (1:2,000, Cell Signaling, #3873S), mouse anti-β-actin (1:2,000, Cell Signaling, #3700S), goat anti-albumin (1:1,000, R&D Systems, #AF3329-SP), or rabbit anti-podocin (1:1,000, Sigma, #P0372). Paraffin sections of kidneys were blocked with 5% donkey or goat serum in PBS-T (0.1% Tween-20) and then stained with rat anti-mouse EMCN (1:200, Abcam, #ab106100), rabbit anti-EGFP (1:300, Sigma, #SAB4701015), mouse anti-CD31 (1:200, Cell Signaling, #3528S), mouse anti-CD34 (1:200, Santa Cruz, #sc-74499), rat anti-CD45 (1:100, Abcam, #ab23910), goat anti-albumin (1:300, R&D Systems, #AF3329-SP), rabbit anti-podocin (1:200, Thermo Fisher, #PA5-79757), and goat anti-smooth muscle actin (1:100, Thermo Fisher, #PA5-18292). Secondary antibodies included donkey anti-rat Alexa Fluor 594 (1:300, Thermo Fisher, #A21209), goat anti-rabbit Alexa Fluor 488 (1:300, Thermo Fisher, #A11008), donkey anti-goat IgG Alexa Fluor 647 (1:300, Thermo Fisher, #A32849), and goat anti-mouse Alexa Fluor 647 (1:300, Thermo Fisher, #A21235).

### Protein extraction and western blot

Freshly collected tissues were stored in Allprotect Tissue Reagent (#76405) or directly lysed in RIPA buffer (Cell Signaling, #9806S) with Protease Inhibitor Cocktail (1:100, Cell Signaling, #5871S) and homogenized (Fisher Scientific, #12-141-361) on ice. After centrifuging at 12,000 g for 5 min, lysate supernatants were prepared with equal protein concentration determined using the Pierce BCA assay kit (Thermo Scientific, #23227). Laemmli’s SDS Sample Buffer (Boston Bio Products, #BP-110R) was added before boiling samples at 95°C for 5 min. Samples were separated by electrophoresis for 2 hr at 60-100 V using 4% to 20% precast gradient gels (MiniPROTEAN TGX; Bio-Rad, #4561094) in 1X SDS-Tris-glycine buffer (Bio-Rad, #1610772EDU). Gradient gels were then transferred to 0.45-μm nitrocellulose membranes (VWR, #27376-991) for 1 hr at 75 V in ice-cold 20% methanol in 1X Tris-glycine buffer (Bio-Rad, #1610771EDU).

Membranes were blocked for 1 hr with 3% BSA in PBS and then probed with primary antibodies for the proteins of interest at 4°C overnight. Corresponding secondary antibodies IRDye® 800CW goat anti-rabbit (Licor, #925-32211) or IRDye® 680RD goat anti-mouse (Licor, #925-68070) were used. Fluorescence LI-COR Odyssey (LI-COR) was utilized for final image development. Band intensities were determined using ImageJ software and normalized to housekeeping proteins.

### Morphologic examination

The mice were subjected to anesthesia via intraperitoneal injection of a ketamine/xylazine mixture (up to 80 mg/kg body weight ketamine and 10 mg/kg body weight xylazine). Subsequently, the rib cage was opened to expose the heart, and a 27G needle was inserted into the left ventricle followed by a cut on the right atrium to initiate exsanguination by PBS perfusion. Kidneys were collected following PBS perfusion and directly fixed in 10% formalin (Fisher Scientific, #23-305510) overnight at 4°C. Fixed kidneys were paraffin-embedded, sectioned (5 μm thick), and stained with hematoxylin and eosin (H&E) to examine glomeruli morphology or with Masson trichrome stain to visualize collagen deposition. Images were acquired with an EVOS FL automated stage live cell imaging system (Life Technologies, Cambridge, MA) or a Zeiss Axioscope 7 (Oberkochen, Germany) and stitched automatically. The number and size of glomeruli were quantified on whole scans of kidney longitudinal section from four sets of mice in a masked manner using Image J. Collagen deposition stained as the blue color in Trichrome staining/total glomeruli area in 63x magnification was quantified in randomly selected images from four sets of mice using Image J in a masked manner.

### Immunohistochemistry

Paraffin sections of kidney were deparaffinized and rehydrated in 100% xylene (3X, 5 min), 100% ethanol (2X, 5 min), 95% ethanol (2X, 5 min), 70% ethanol (2X, 5 min) and distilled water (1X, 5 min). Antigen retrieval was performed by heating the slides (95°C to 100°C) for 25 mins in 10 mM sodium citrate buffer (pH 6.0) and cooling at room temperature (RT) for 30 min. Sections were then washed with distilled water, permeabilized with 0.5% Triton X-100 in PBS for 5 mins, and blocked (5% goat or donkey serum in PBS) for one hr at RT. The primary antibody was prepared in blocking buffer, and slides were incubated with the primary antibodies overnight at 4°C in a humidified chamber. Sections were washed with PBS and incubated with secondary antibodies in blocking buffer for two hr at RT. Slides were mounted using Prolong Gold Antifade Reagent with DAPI (Invitrogen, #P36935). Images were obtained using Zeiss Axioscope 7 (Oberkochen, Germany) or Leica TCS SP8 Confocal Microscopy (Leica Microsystems, Wetzlar, Germany). CD45^+^ cells were quantified on 63X images from randomly selected 17 images (each genotype) from four sets of mice using the cell counting function on ImageJ in a masked manner.

### Flow cytometry

Flow cytometry was performed following previously described methods^21,22^. Briefly, mice (20 to 24 week-old) were euthanized, and fresh kidneys were harvested and minced into small pieces. The tissue was then digested with an enzymatic solution consisting of 60 U/mL DNase I (Sigma-Aldrich, #D4513) and 450 U/mL collagenase type IV (Gibco, 17104019) in 20 mM HEPES (Gibco, #15630106) at 37°C for one hr. After passing the digested tissue through a 70 µm filter, the cells were resuspended in Flow Cytometry Staining Buffer (FACS buffer, eBioscience, #00-4222-57), and incubated with Fc-blocking antibody (1:100, eBioscience, #14-9161-73) for 30 min at RT. Next, the cells were stained with the following antibodies: FITC anti-CD45 (#157608), PerCP anti-CD11b (#101230), APC anti-CD3 (#100236), PE/Cy5 anti-CD19 (#115510), APC/Cy7 anti-Ly6C (#128016), Pacific blue anti-Ly6G (#127612), PE-conjugated CD144 (#138009), and PE/Cy7-conjugated CD4 (#100422) (all obtained from Biolegend and used at a 1:100 dilution in 100 µL blocking buffer). After two washes with FACS buffer, the cells were evaluated using a BD LSR II Analyzer (BD Biosciences) and plotted using FlowJo V10.8 software (Tree Star).

### Urine collection and analysis

Spot urine samples were collected from mice at three different times per day (10 am, 1 pm, and 4 pm) and combined for analysis. Equal volumes of urine samples (2 µL for Coomassie blue staining and 1 µL for western blot) were loaded onto precast 4-20% gradient gels for electrophoresis. After the electrophoresis separation of urine samples, the gels were rinsed in PBS and incubated with Coomassie Brilliant Blue R-250 (BioRad, #161-0436) for 2 hr at RT. The gels were then destained overnight on a shaker using a freshly prepared destaining buffer (10% acetic acid, 50% methanol, and 40% distilled H_2_O).

To quantify albumin concentrations in urine, a mouse albumin ELISA assay (Crystal Chem, #80630) was performed according to the manufacturer’s instructions. In brief, 5 µL of urine or standard concentrations of mouse albumin in a total volume of 100 µL PBS were added to a plate and incubated for 30 min at RT. After washing, a 100 µL aliquot of albumin antibody conjugate was added and incubated for another 30 min at RT. Following a 10-minute incubation with 100 µL of substrate solution, 100 µL of stop solution was added, and the optical density at 450/630 nm was measured.

Urine creatinine concentrations were determined using an enzymatic-based mouse creatinine Assay Kit (Crystal Chem, #80350). Eight microliters of urine samples were mixed with 270 µL of sarcosine oxidase solution and incubated for 5 min at 37°C. The optical density at 550 nm was measured, and then 90 µL of peroxidase solution was added. After another 5-min incubation at 37°C, the optical density at 550 nm was measured for quantification.

### Transmission electron microscopy

Mice were perfused as described above through the left cardiac ventricle with PBS and followed by half Karnovsky’s fixative (2% formaldehyde + 2.5% glutaraldehyde, in 0.1 mol/L sodium cacodylate buffer, pH 7.4) for perfused fixation of tissues. Kidneys were then collected and immersed in half Karnovsky’s fixative for 24 hr at 4°C. After fixation, samples were processed by the Morphology Core at Schepens Eye Research Institute of Mass. Eye and Ear for TEM imaging.

In brief, the kidneys were trimmed into 1-mm thick segments and rinsed with 0.1 mol/L sodium cacodylate buffer. They were then post-fixed and en bloc stained with 2% gadolinium triacetate in 0.05 mol/L sodium maleate buffer (pH 6) for 30 min. Following dehydration and polymerization of the samples using a 60°C oven, semithin sections (1 μm thickness) were cut and stained with 1% toluidine blue in a 1% sodium tetraborate aqueous solution for examination under light microscopy. Ultrathin sections were obtained from the grids and stained with aqueous 2.5% gadolinium triacetate and modified Sato lead citrate. Grids were imaged using an FEI Tecnai G2 Spirit transmission electron microscope (FEI, Hillsboro, OR) at 80 kV interfaced with an AMT XR41 digital charge-coupled device camera (Advanced Microscopy Techniques, Woburn, MA) for digital image acquisition at 2000 × 2000 pixels at 16-bit resolution. Width and length of podocyte foot processes as well as the thickness of the glomerular basement membrane were quantified on 25,000x TEM images from 23 (each genotype) randomly selected images from 3 sets of mice using the measurement function on ImageJ in a masked manner.

### Blood pressure measurement

The CODA non-invasive blood pressure system (Kent Scientific, Torrington, CT) was used to measure tail blood pressure in mice from the tail as previously described^23^. CODA system was factory calibrated prior to experiments. Standard settings and recommendations were used as follows. The mice were acclimated for five to seven consecutive days before baseline blood pressure measurements were taken. Mice were acclimated for 1 hr in a quiet room (22 ± 2°C) before measurements and then gently restrained in tubes with adjustable end holders to minimize movement. The occlusion cuff was positioned at the tail base, with the VPR sensor cuff adjacent. Heating pads warmed the tails to 33-35°C for 5 min before and during recordings. For each measurement cycle, the occlusion cuff inflated to 250 mmHg then slowly deflated over 20 seconds while the VPR sensor detected tail volume changes as blood flow returned. A 15 μl minimum volume change threshold was set. Each recording session consisted of 15-20 cycles and the first five acclimation cycles were excluded. The blood pressure of both male and female mice was measured but analyzed separately. Systolic, diastolic and average blood pressure were quantified.

### Fluorescence activated cell sorting (FACS) for endothelial cells and podocyte isolation

The method for glomeruli isolation for FACS sorting was adapted from previously published protocols^24-26^. The mice (female littermates at 6-month-old) were quickly perfused with cold PBS before kidneys were harvested and decapsulated. The two kidneys from one mouse were then minced into 1-2 mm pieces on ice and combined as one sample then digested in digestion buffer I (Collagenase I at 1.6 mg/ml and DNase I at 7.5 ug/ml in HBBS with Ca^2+^ and Mg^2+^) at 37°C for 40 min with gentle agitation. The digested tissue was neutralized with FBS and passed through a 100 μm cell strainer then washed once with 5-10 ml wash buffer (1 mM sodium pyruvate, 1 x MEM nonessential amino acids, 5 mM L-glutamine, 25 mM HEPES in DMEM). The flow-through was filtered through a 40 μm cell strainer. Undigested glomeruli that remain in the strainer were washed with 2-5 ml of wash buffer and glomeruli collected by washing the interior of the strainers with cold HBBS for at least 10 times. The glomeruli were pelleted, then resuspended and digested in prewarmed digestion buffer (1mg/ml Liberase TL, 1mg/ml collagenase IV and 50 ug/ml DNase I in DMEM) for 60 min at 37°C. After neutralization with FBS, the cell suspension was centrifuged and resuspended in FACS buffer (1% FBS in HBBS) with Fc block (1:100) for 15 min at RT then incubated with PE-CD31 (#102407), APC podoplanin (#127409), BV785 CD45 (#103149) antibodies in blocking buffer at RT for 30 min (all from Biolegend and used at a 1:100 dilution). Cells were DAPI stained before sorting through BD FACSAria IIu Cell Sorter. Forward and side scatter gates were set to exclude debris and cell doublets/aggregates. The CD31^+^ and CD45 ^-^ populations were identified from the DAPI^-^ cells and isolated as the endothelial fraction. The podoplanin^+^ cells from CD31/CD45 double-negative cells were isolated as the podocyte fraction. Sorted ECs and podocytes were collected in 10% FBS in DMEM with RNase inhibitor (VWR, # N2611) and centrifuged and resuspended in Trizol for RNA extraction.

### RNA seq of sorted ECs and podocytes and data analysis

Total RNA was extracted from sorted endothelial cells and podocytes using a QIAGEN RNeasy Micro Kit with on column DNase digestion according to the manufacturer’s instructions. Samples were submitted for ultra-low input RNA-seq to GENEWIZ from Azenta Life Sciences (South Plainfield, NJ, USA). The cDNA libraries were generated with the Ultra Low Input RNA Library Prep Kit (Takara Bio) and then sequenced on an Illumina 6000 using paired-end sequencing with a read length of 30 million base pairs. The raw FASTQ files was provided by Genewiz for downstream bioinformatics analysis. Reads were trimmed to remove adapter sequences using Trim Galore! (Version 0.6.5). Quality control on both raw reads and adaptor-trimmed reads was performed with MultiQC. Reads were aligned to the mm10 genome using Rsubread v1.5.327^27^.

Gene counts were quantified by Entrez Gene IDs using featureCounts and Rsubread’s built-in annotation^27^. Gene symbols were from NCBI gene annotation. Genes with count-per-million above 0.5 in at least three samples were retained in the analysis. Differential expression analysis was performed using limma-voom and DESeq2 in R28. Genes with log2 fold change >0.5 and p value <0.01 were considered to have significant differences of expression. The Database for Annotation, Visualization and Integrated Discovery (DAVID 6.8) and ClusterProfiler package in R were used to identify functional categories for the genes enriched and genes with significant differences of expression29. Gene Ontology Biological Process (GO_BP) with use of all 313 genes from endothelial cells and 216 genes from podocytes protein-coding genes with significant differences of expression. Data discussed in this publication have been deposited in NCBI’s Gene Expression Omnibus and are accessible through GEO Series accession number GSE271432.

### Statistical analysis

All values are expressed as mean ± SEM. Statistical analysis was performed using an unpaired Student’s t-test or one-way-ANOVA with Tukey post hoc test for multiple comparisons (GraphPad Prism 9, GraphPad Software Inc., San Diego, CA, USA). A p value <0.05 was considered statistically significant. Each experimental condition included at least three technological replicates, and all experiments were independently repeated at least three times.

## RESULTS

### Validation of EMCN deletion

To determine the functional role of EMCN in vivo, we used the Cre-lox system to generate EMCN KO mice, as shown in Figure 1A. The global EMCN KO mice are viable and fertile. To confirm successful deletion in EMCN^+/-^ and EMCN^-/-^ mice, tissues including the kidney, lung, thyroid, spleen, small intestine, and liver were collected, and EMCN mRNA levels were examined (Figure 1B). EMCN mRNA levels in EMCN^+/-^ tissues were significantly reduced and were not detectable in EMCN^-/-^ tissues compared to tissues from EMCN^+/+^ mice. Levels of EMCN protein were also examined in kidneys, lungs, and thyroids and consistent with the mRNA levels, EMCN protein expression was significantly reduced in tissues from EMCN^+/-^ mice and completely absent in tissues from EMCN^-/-^ mice (Figure 1C).

To examine the expression pattern of EMCN in kidneys, immunohistochemistry was performed on paraffin sections of kidneys from EMCN^+/+^, EMCN^+/-^, and EMCN^-/-^ mice (Figure 1D). EMCN was only detected in kidneys of EMCN^+/+^ and EMCN^+/-^ mice where it is primarily expressed in the glomeruli and co-localized with the endothelial marker CD31. EGFP, a reporter of Emcn gene deletion was detected in the glomeruli and co-localized with CD31 in EMCN^+/-^ and EMCN^-/-^ mice, confirming the endothelial-specific expression of EMCN. EMCN expression was absent in EMCN^-/-^ kidneys.

### Glomeruli of EMCN^-/-^ mice displayed increased CD45^+^ cells infiltration

Gross morphological examination of the EMCN^-/-^ mice and their organs suggested no obvious differences compared to the wildtype littermates. No significant differences in body weights were found among male (Supplemental Figure 2A) and female (Supplemental Figure 2B) mice of the three genotypes at 3 weeks to 6 months old. Because we have reported a role for EMCN in preventing endothelial-leukocyte adhesion^14^, we examined infiltration of leukocyte in the kidney of the EMCN^-/-^ mice to determine tissue inflammatory status in the absence of this anti-adhesion barrier for leukocytes. Immunohistochemical analysis revealed an increased number of CD45^+^ cells in the glomeruli of EMCN^-/-^ mice compared to EMCN^+/+^ and EMCN^+/-^ mice (Figure 2A). Quantification of the average CD45^+^ cells per kidney section area revealed 56.17 ± 5.086, 52.06 ± 4.62, and 94.90 ± 9.902 CD45^+^ cells per mm^2^ in the EMCN^+/+^, EMCN^+/-^ and EMCN^-/-^ kidney sections respectively, confirming a significant increase in CD45^+^ leukocyte infiltration in the EMCN^-/-^ kidney compared to heterozygous (*p* < 0.0001) and wildtype littermates (*p* < 0.001) (Figure 2B).

**Figure 2.**
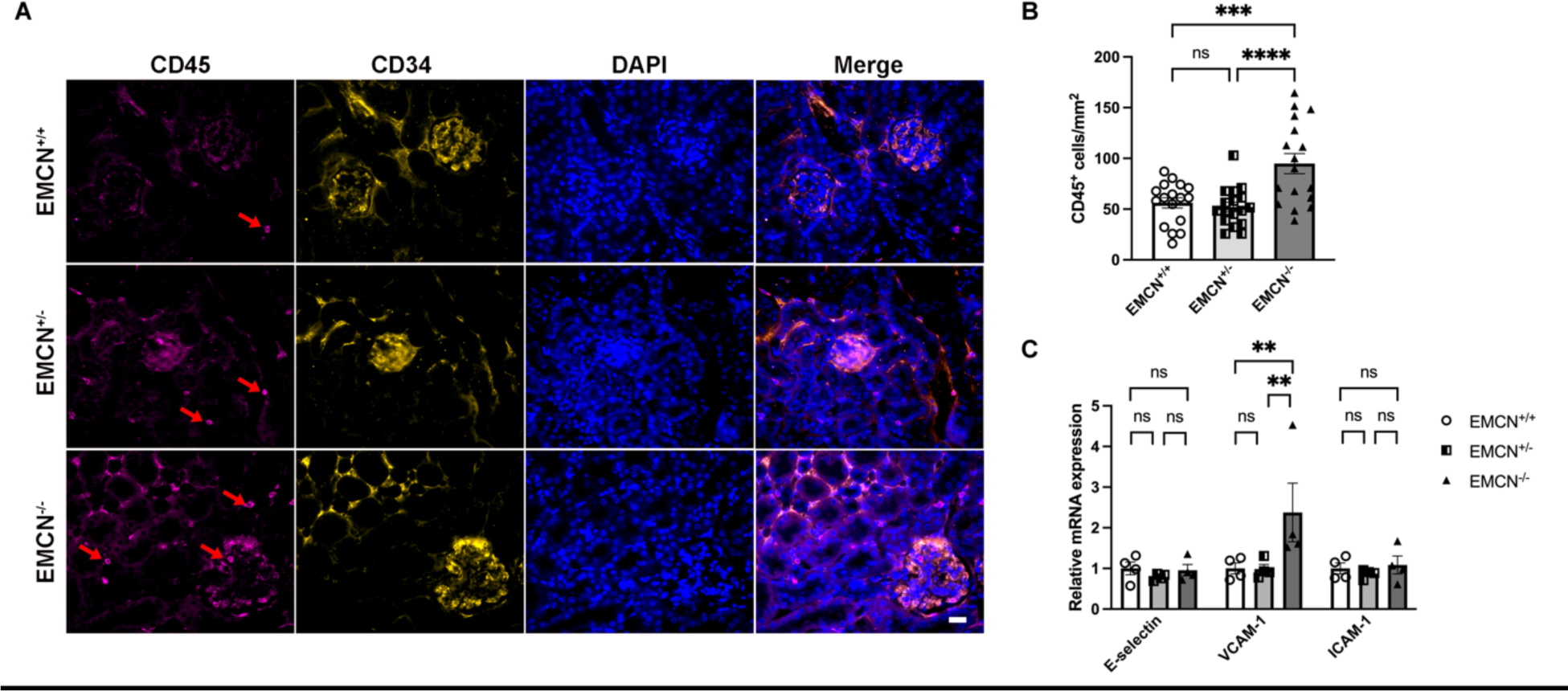
Increased CD45^+^ cell infiltration in kidney of EMCN^-/-^ mice. **(A)** Kidneys from EMCN^+/+^, EMCN^+/-^, and EMCN^-/-^ mice (16 to 24 weeks old) were collected and fixed with 10% formalin overnight for paraffin sectioning. Immunohistochemical staining was performed to visualize CD45 and CD34; nuclei were stained with DAPI. n = 4. Scale bar = 20 μm. **(B)** Quantification of CD45^+^ cells in kidney sections. n = 17 images from four sets of EMCN^+/+^, EMCN^+/-^, and EMCN^-/-^ sections. *** *p* < 0.001, **** *p* < 0.0001, one-way ANOVA. **(C)** mRNA from kidneys from EMCN^+/+^, EMCN^+/-^, and EMCN^-/-^ mice was extracted and examined for E-selectin, VCAM-1, and ICAM-1 expression. n = 4, ** *p* < 0.005, one-way ANOVA.

Because endothelial adhesion molecule expression can directly affect leukocyte attachment and infiltration, mRNA expression levels of E-selectin, vascular cell adhesion molecule 1 (VCAM-1), and ICAM-1 were also examined. A significant increase in VCAM-1 mRNA expression in the kidney was detected in EMCN^-/-^ mice when compared to both EMCN^+/+^ and EMCN^+/-^ littermates (1.47 ± 0.454 vs. 1.00 ± 0.008 and 0.88 ± 0.055, *p* < 0.005 for both comparisons) (Figure 2C). No significant differences were observed in mRNA levels for E-selection and ICAM-1 among the three genotypes. Since no significant differences were detected between EMCN^+/+^ and EMCN^+/-^ in CD45^+^ cell infiltration and adhesion molecule expression, subsequent studies were focused on EMCN^+/+^ and EMCN^-/-^ mice. To further explore the identity of the infiltrated CD45^+^ cells, fresh kidneys were collected, and single-cell suspensions were prepared for flow cytometry analysis. The single cell suspensions were first sorted for leukocyte common antigen CD45, then the CD45-positive population was sorted for B-lymphocyte antigen CD19, the T-lymphocyte marker CD3, or the myeloid cell lineage marker CD11b. The CD45^+^ CD11b^+^ population was then sorted for Ly6C ^high^ Ly6G ^high^ and Ly6C ^high^ Ly6G ^low^ groups, which represent neutrophils and monocyte/macrophages, respectively. The number of ECs in kidney cell suspension was assessed using CD144 for sorting (Figure 3A). Consistent with the immunohistochemical observations, there was an increase in the proportion of CD45^+^ cells (Figure 3B) in EMCN^-/-^ compared to EMCN^+/+^ mice (11.4 ± 0.151% vs 7.5 ± 0.292%, n = 10, *p* < 0.05, Student’s t-test). Among the CD45^+^ cells, there was a significant increase in cells of the monocytes/macrophage lineage (Figure 3D), marked by Ly6C^high^ Ly6G^low^ in EMCN^-/-^ compared to EMCN^+/+^ kidneys (1.60 ± 0.254% vs 1.09 ± 0.508%, n = 10, *p* < 0.05, Student’s t-test). The percentage of neutrophils, T lymphocytes, B lymphocytes, and ECs (Figure 3 C, E, F, G) in the kidney was not significantly changed between EMCN^+/+^ and EMCN^-/-^ mice (1.00 ± 0.003% vs 1.04 ± 0.230%, 0.07 ± 0.021% vs 0.09 ± 0.019%, 0.079 ± 0.029% vs 0.107 ± 0.019%, 2.970 ± 0.018% vs 3.150 ± 0.535%, *p* < 0.05 for all, Student’s t-test).

**Figure 3.**
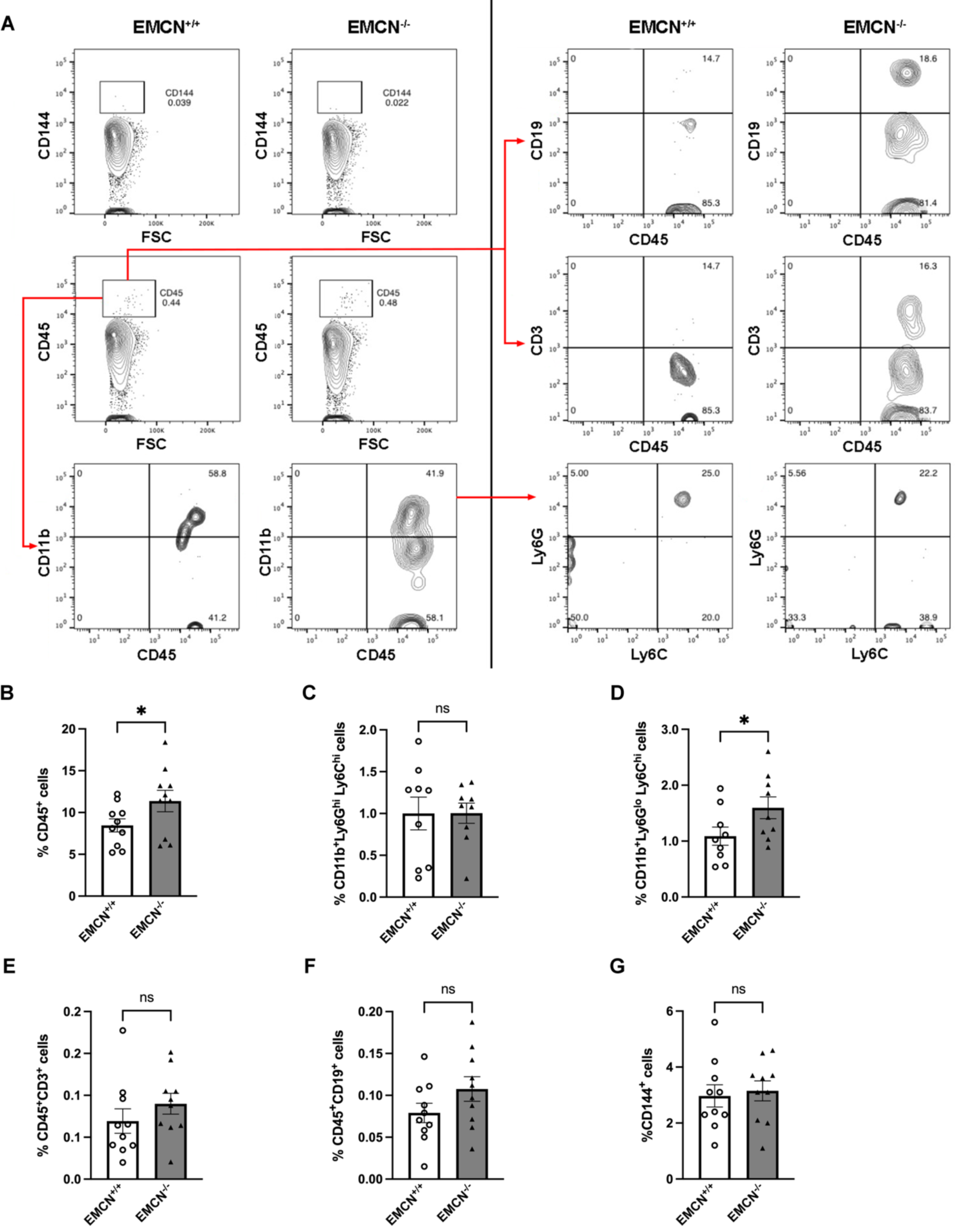
Increased CD11b^+^Ly6G^lo^Ly6C^hi^ cells in kidneys of EMCN^-/-^ mice. **(A**) Representative images of flow cytometry of kidney cells isolated from EMCN^+/+^ and EMCN^-/-^ mice (16 to 24 weeks old). Percentage of **(B)** CD45^+^ cells, (**C)** CD11b^+^Ly6G^hi^Ly6C^hi^ cells, **(D)** CD11b^+^Ly6G^lo^Ly6C^hi^ cells, **(E)** CD45^+^CD3^+^ cells, and **(F)** CD45^+^CD19^+^, (G) CD144^+^ ECs. n = 10, * *p* < 0.05, Student’s t-test.

### EMCN^-/-^ kidneys display smaller glomeruli and increased collagen deposition

We next examined the morphology of the kidneys and found that there was no significant difference in gross morphology, kidney size, and weight between EMCN^+/+^ and EMCN^-/-^ mice (Figures 4A-B). Quantification of glomeruli number and glomeruli area from H&E sections of kidneys revealed a similar number of glomeruli between EMCN^+/+^ and EMCN^-/-^ mice (173.5 ± 16.54 vs 180.0 ± 16.42, *p* < 0.05, Student’s t-test), however, the average size of glomeruli in EMCN^-/-^ was smaller when compared to EMCN^+/+^ mice (3646 ± 279.0 pixels vs. 4363 ± 185.3 pixels, *p* < 0.05, Student’s t-test) (Figures 4C-E). Trichrome staining for the ECM of kidney sections demonstrated increased collagen deposition per glomeruli area in EMCN^-/-^ kidney sections compared to that from EMCN^+/+^ (Figure 4C, F) (101.9 ± 16.54 5.455 pixels/um^2^ vs 84.33 ± 4.822 pixels/um^2^, *p* < 0.05, Student’s t-test).

**Figure 4.**
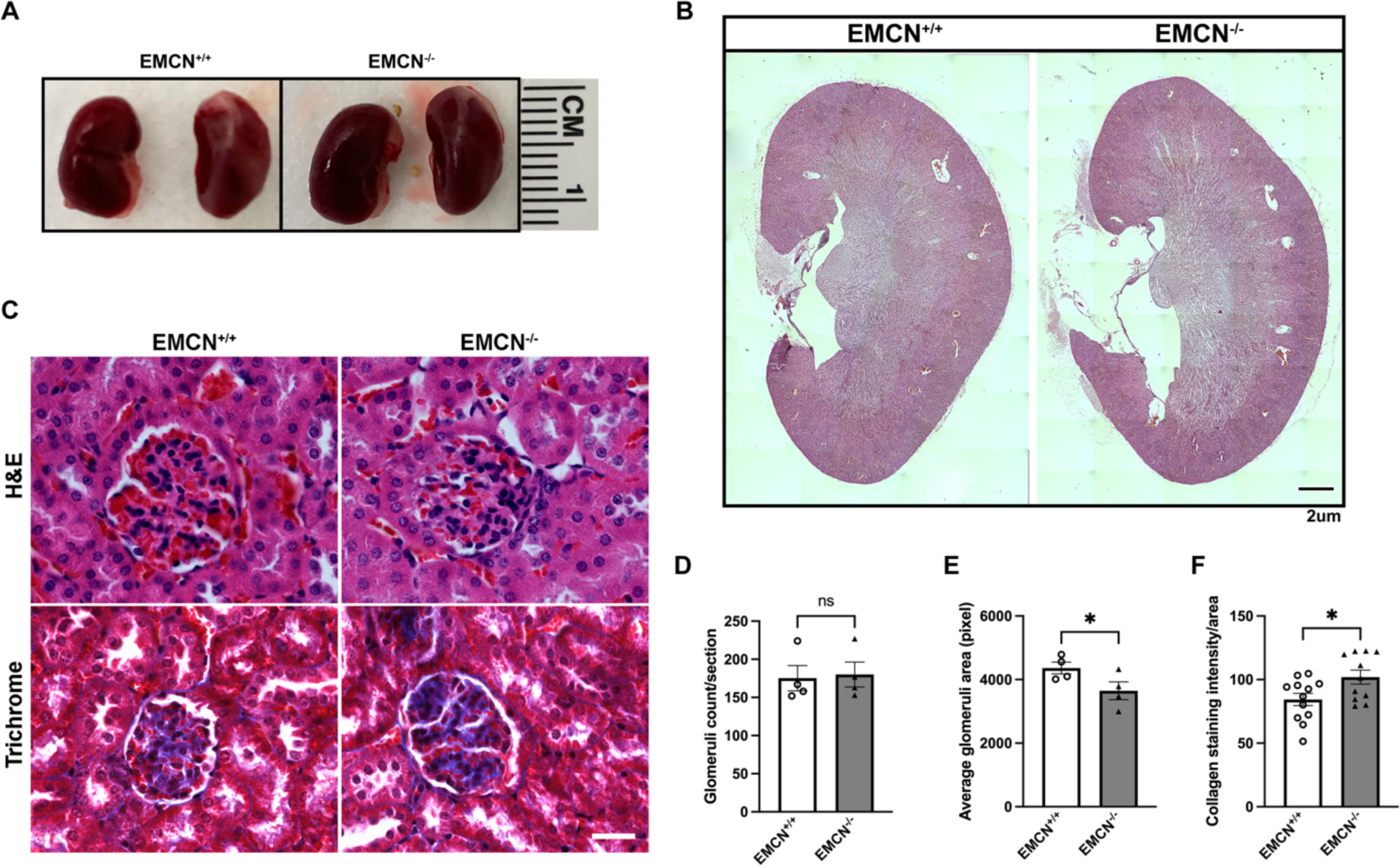
Histology of kidneys from EMCN^+/+^ and EMCN^-/-^ mice, and increased collagen staining in kidneys from EMCN^-/-^ mice. **(A, B)** Representative gross morphology images of kidneys and H&E-stained kidney coronal sections across the renal pelvis from EMCN^+/+^ and EMCN^-/-^ mice (16 to 24 weeks old). n = 3, Scale bar = 1 mm. (**C)** Representative images of kidney sections with H&E staining (upper panel) and Trichrome staining (lower panel) from EMCN^+/+^ and EMCN^-/-^ mice for morphometric analysis. n = 3, Scale bar = 20 μm. **(D)** Glomeruli number and **(E)** glomeruli areas were quantified using ImageJ on H&E stained-kidney sections from EMCN^+/+^ and EMCN^-/-^ mice. n = 4, * *p* < 0.05, Student’s t-test. **(F)** Collagen staining intensity (blue color from Trichrome staining in C) per glomeruli area was quantified using ImageJ. n = 12 sections from four animals, * *p* < 0.05, Student’s t-test.

### EMCN^-/-^ mice exhibit impaired glomerular barrier function with albuminuria

Because of the reduction in size and increased extracellular matrix (ECM) deposition seen for the glomeruli in the absence of EMCN, we next investigated if there were any functional changes in the kidneys in EMCN^-/-^ mice. Coomassie Blue staining of urine from EMCN^+/+^ and EMCN^-/-^ mice was shown in Figure 5A. While there were similar levels of major urinary proteins (MUPs; observed around 16 kDa), a band of the molecular weight corresponding to albumin (around 60 kDa) was significantly higher in urine collected from EMCN^-/-^ mice compared to EMCN^+/+^ mice. Total urine creatinine, albumin, and total protein concentrations were quantified using a combination of Coomassie Blue staining, ELISA, and enzymatic-based assays. While no significant change in urine creatinine concentration was observed between EMCN^+/+^ and EMCN^-/-^ (68.64 ± 4.877 mg/dl vs. 68.28 ± 1.838 mg/dl, *p* < 0.05, Student’s t-test) (Figure 5B), there was a trend of increased total protein/creatinine in EMCN^-/-^ mice compared to EMCN^+/+^ mice (71.23 ± 4.946 g/g vs. 56.65 ± 7.258 g/g, *p* > 0.05, Student’s t-test) (Figure 5C). A significant increase in urine albumin/creatinine was detected in EMCN^-/-^ mice compared to EMCN^+/+^ mice (47.71 ± 13.76 ug/g vs. 14.15 ± 2.327 ug/g, *p* < 0.05, Student’s t-test) (Figure 5D). To confirm the increased urine albumin in EMCN^-/-^ mice, fresh urine samples were collected from both EMCN^+/+^ and EMCN^-/-^ mice and examined by western blot analysis (Figure 5E). The quantification of albumin band intensity was consistent with our ELISA data and confirmed increased urine albumin in EMCN^-/-^ mice compared to EMCN^+/+^ mice (25642 ± 6722 pixels vs. 11646 ± 2724 pixels, *p* < 0.05, Student’s t-test) (Figure 5F). To investigate the localization of the increased albumin, immunohistochemistry for albumin, EMCN, and CD34 was performed on kidney sections (Figure 5G). In the glomeruli of EMCN^+/+^ mice, ECs are marked by EMCN and CD34, and some albumin was visualized inside the capillary lumen. In contrast, in the glomeruli of EMCN^-/-^ mice, increased intensity of albumin staining was observed in the Bowman’s space in the kidney. Given the observed increase in albuminuria, we investigated whether alterations in blood pressure in EMCN^-/-^ mice could contribute to the increased albuminuria levels. Alternatively, the increased albuminuria could be due to structural and functional changes in the glomerular filtration barrier (GFB) or a combination of both hemodynamic and filtration barrier defects. Evaluation of systolic (male: 131.2 ± 4.9 mmHg vs 124.9 ± 2.9 mmHg; female: 122.9 ± 3.8 mmHg vs 123.1 ± 3.3 mmHg), diastolic (male: 105.8 ± 4.8 mmHg vs 97.7 ± 3.4 mmHg; female: 96.9 ± 3.6 mmHg vs 99.5 ± 3.0 mmHg), and average arterial blood pressure (male: 113.8 ± 4.8 mmHg vs 105.9 ± 3.4 mmHg; female: 105.1 ± 3.7 mmHg vs 106.7 ± 3.2 mmHg) revealed no significant differences between EMCN^-/-^ mice and their EMCN^+/+^ littermate controls in either males or females (Figure 6AB). Power analysis calculations with 80% power and a p-value < 0.05, based on the previously published data set^28^, indicated that a sample size of 4 mice per group would be sufficient to detect significant differences. In our blood pressure measurements, we included more than 6 mice in each group, which provided adequate power to detect differences in blood pressure changes in EMCN^-/-^ compared to controls.

**Figure 5.**
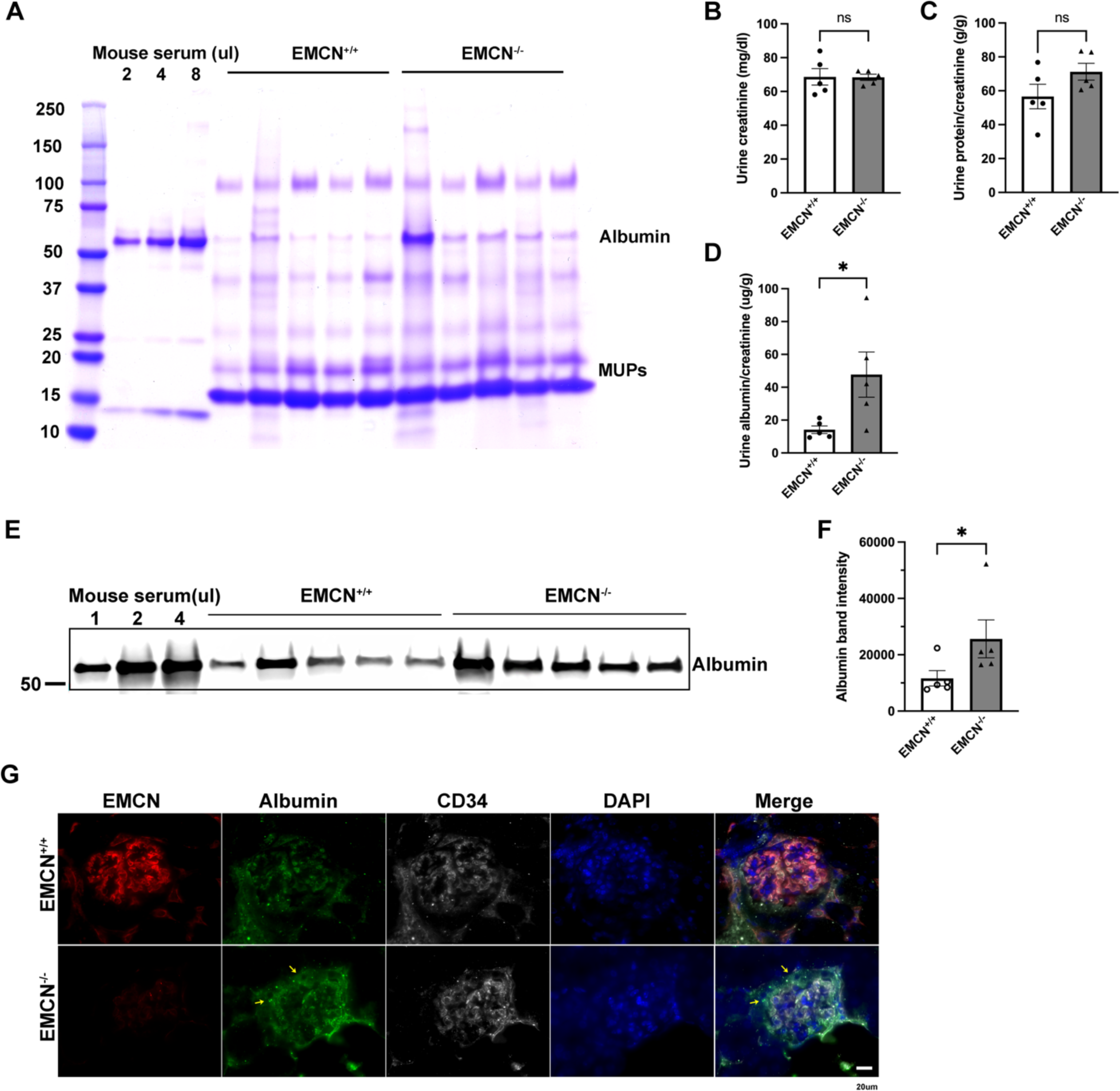
Increased albumin in urine and Bowman’s space in kidneys of EMCN^-/-^ mice. Urine samples were collected from 16 to 24-week-old EMCN^+/+^ and EMCN^-/-^ mice as described in the methods. **(A)** Two µl of each urine sample were loaded on SDS-page gel with mouse serum as a control and stained with Coomassie blue. Putative albumin band and MUPs bands were indicated as shown. **(B-D)** Mouse urine creatinine and albumin concentrations were quantified by ELISA and urine creatinine concentration, urine protein/creatinine, and urine albumin/creatinine levels from EMCN^+/+^ and EMCN^-/-^ mice were compared. n = 5, * *p* < 0.05, Student’s t-test. **(E)** Representative image of western blot examining the albumin expression level in one µl of freshly collected urine samples from EMCN^+/+^ and EMCN^-/-^ mice. **(F)** Quantification of albumin band intensity on western blot of urine samples from EMCN^+/+^ and EMCN^-/-^ mice. n = 5, * *p* < 0.05, Student’s t-test. **(G)** Representative images of immunohistochemical staining to visualize the expression of EMCN, albumin, and CD34 in kidney sections from EMCN^+/+^ and EMCN^-/-^ mice; nuclei were stained with DAPI. n = 4, Scale bar = 20 μm.

**Figure 6.**
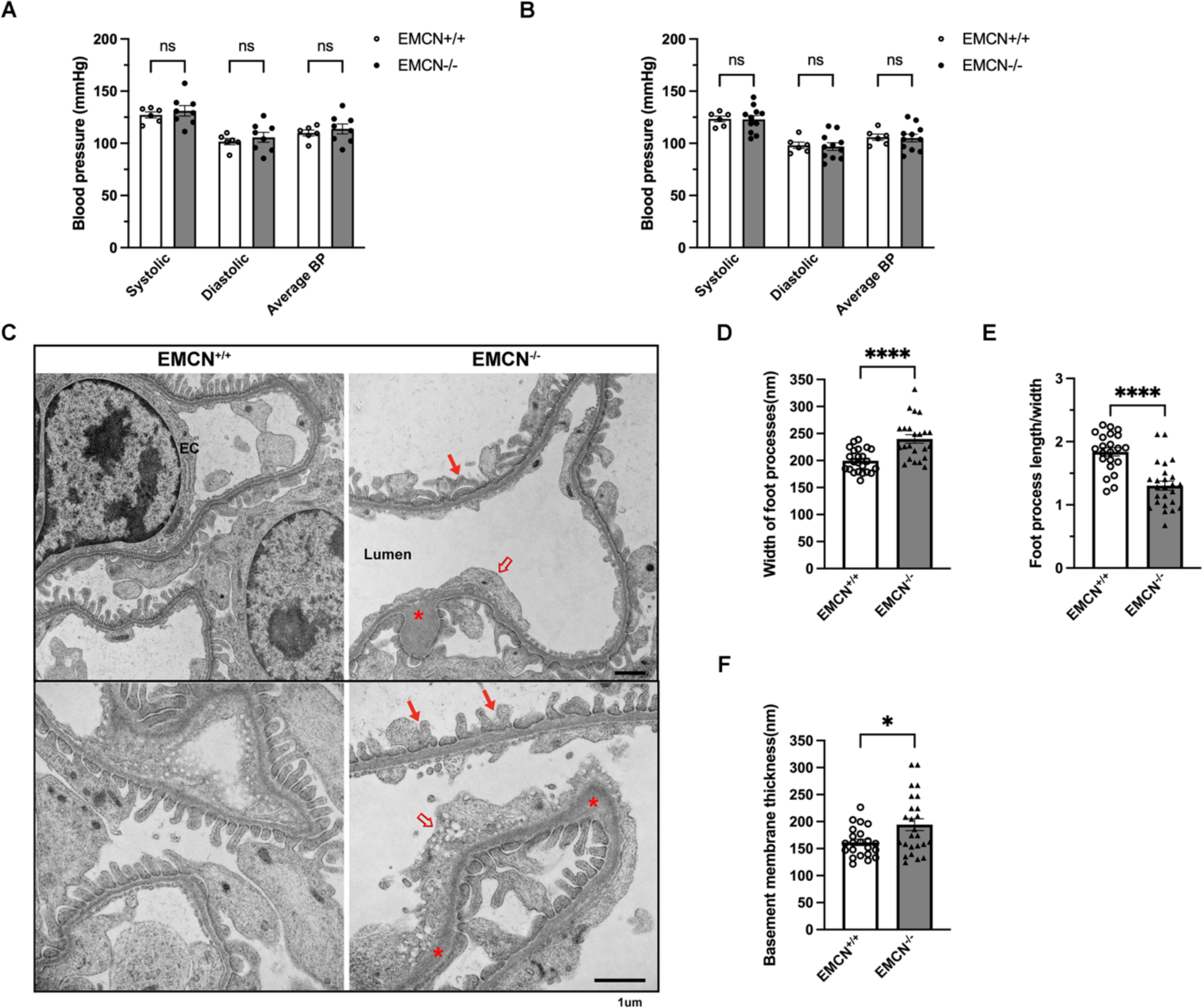
Defective glomerular ultrastructure in kidneys from EMCN^-/-^ mice. **(A)** Systolic, diastolic, and average blood pressure measured by a non-invasive tail-cuff blood pressure system in female littermates and **(B)** male littermates. n = 6, 8 for female, n = 6, 11 for male. *p* > 0.05, Welch’s t-test. **(C)** Representative TEM images of the GFB (fenestrated endothelium, basement membrane, and podocytes) showing lower magnification (upper panels) and higher magnification (lower panels) from EMCN^+/+^ (left panels) and EMCN^-/-^ (right panels) mice. Red arrows point to the effaced and fused podocytes, red hollow arrows point to disorganized fenestration of the endothelium, and red asterisks mark the thickening of basement membrane in the EMCN^-/-^ mice. n = 4, Scale bar = 1 μm. **(D)** Width of podocyte foot processes, **(E)** length/width ratio of podocyte foot process, and **(F)** basement membrane thickness were quantified in a masked fashion in three sets of EMCN^+/+^ and EMCN^-/-^ samples by ImageJ software. n = 23, * *p* < 0.05, **** *p* < 0.0001, Student’s t-test.

### Disrupted ultrastructure of the GFB in kidneys of EMCN^-/-^ mice

We then examined the ultrastructure of EMCN^-/-^ kidneys using TEM. Representative TEM images of glomeruli from EMCN^+/+^ kidneys show the well-structured and organized endothelial fenestrations, uniform thickness of the basement membrane, and the regularly shaped foot processes of podocytes, which together form the functional podocyte filtration slits and GFB (Figure 6C, left panels). In contrast, glomeruli of EMCN^-/-^ kidneys exhibit thickened glomerular ECs and disorganized fenestrations, regionally thickened basement membrane, and podocyte foot processes that are effaced and fused and together exhibit a defective GFB (Figure 6C, right panels). Masked quantification of TEM images indicated that effaced podocyte foot process and thickened GBM were observed in EMCN^-/-^ mice (Figure 6 D-F). The podocyte foot processes in EMCN^-/-^ mice were significantly wider than in wildtype controls (239.9 ± 7.877 nm vs. 199.7 ± 4.466 nm, *p* < 0.0001, Student’s t-test), displayed decreased foot process length (length/width ratio, 0.675 ± 0.072 vs. 1.211 ± 0.065, *p* < 0.0001, Student’s t-test), and increased basement membrane thickness (194.3 ± 10.90 nm vs 161.5 ± 5.83 nm, *p* < 0.05, Student’s t-test) compared to that from EMCN^+/+^ littermates. We also examined the ultrastructure of GFB in aged EMCN^+/+^ and EMCN^-/-^ (21-24 months old) and found similar but more severe changes, specifically in basement membrane thickening and podocyte foot process fusion and effacement, suggesting a progressive degeneration of the GFB in the EMCN^-/-^ kidneys (Supplementary Figure 3).

### Loss of EMCN leads to gene expression changes in glomerular endothelial cells that are involved vascular homeostasis

To gain deeper insights into the molecular mechanisms underlying the structural and functional GFB defects in EMCN^-/-^ kidneys, we isolated glomeruli from freshly harvested kidneys and used FACS to purify glomerular ECs and podocytes for RNA-seq (Figure 7A). As shown in Figure 7B, DAPI negative live cells were gated for CD45^-^ CD31^+^ endothelium population and CD31^-^ podoplanin^+^ podocyte. For each of the sorted endothelial and podocyte populations, between 30,000 and 100,000 cells were collected for ultra-low input RNA-seq. All samples passed quality control checks following library preparation and initial sequencing data analysis.

**Figure 7.**
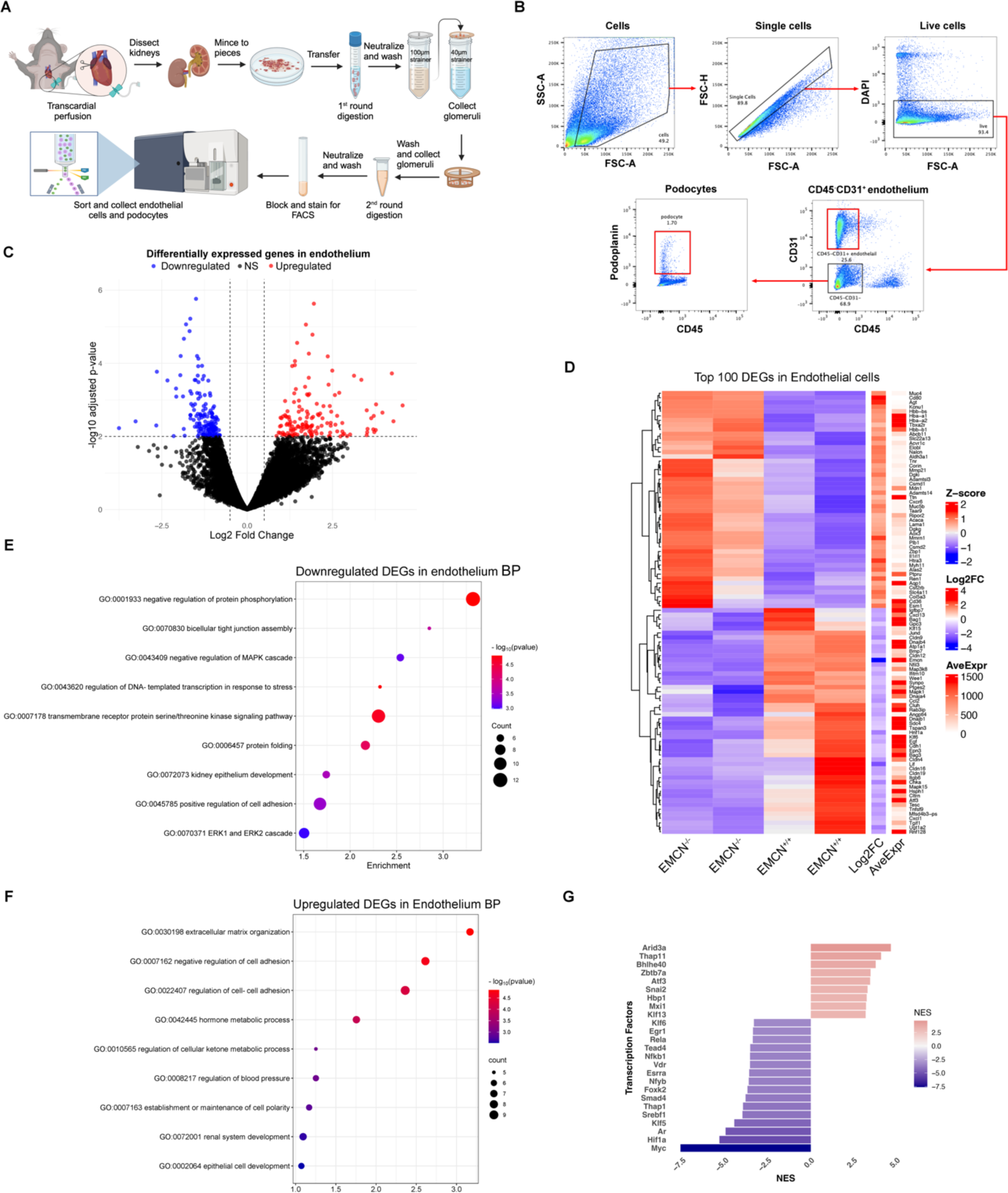
Kidney ECs RNA-seq and differential expressed genes analysis. **(A)** Schematic of the workflow for glomeruli isolation from kidney followed by FACS sorting of EC and podocyte populations for downstream RNA sequencing. n = 2. **(B)** Representative flow cytometry plot illustrating the gating strategy employed to identify and sort live ECs and podocytes. **(C)** Volcano map of differentially expressed genes in ECs, except EMCN, in EMCN^-/-^ kidney endothelium compared to EMCN^+/+^. *p* < 0.01 with log2FC > 0.5 is considered as significantly expressed genes. Blue and red dots indicated downregulated and upregulated genes in EMCN^-/-^ kidney endothelium respectively. **(D)** Heatmap of the top 100 DEGs with base mean > 50 in EMCN^-/-^ kidney endothelium compared to EMCN^+/+^. **(E)** Gene ontology biological process analysis of downregulated genes and **(F)** upregulated genes in EMCN^-/-^ kidney endothelium compared to EMCN^+/+^. **(G)** DoRothEA regulons analysis of transcription factor from DEGs in EMCN^-/-^ kidney endothelium compared to EMCN^+/+^.

A total of 313 differentially expressed genes (DEGs) were found in EMCN^-/-^ glomeruli endothelium compared to EMCN^+/+^ controls, as shown in the volcano plot (Figure 7C). Of which, 157 genes were downregulated, while 156 genes showed upregulated expression. The heatmap of top DEGs (Figure 7D) and the GO analysis of biological processes (Figure 7E) revealed the downregulated genes pathways involved in protein phosphorylation, MAPK cascade, and ERK cascade like Mapk1, Map3k8, Mapk15, Dusp10, Bmp7, Hsph1, Atf3, Errfi1, Ccl2, and Ccn1. Another distinct category of downregulated pathways includes genes associated with tight junction assembly and organization, such as Cdh1, Cldn4, Cldn19, Cldn16, and Cldn9. Additionally, other glycocalyx components such as Gpc3 and Sdc4 were also found significantly downregulated. We identified several genes that were significantly upregulated in the heatmap of top DEGs (Figure 7D) and by GO analysis of biological processes (Figure 7F). Notably, genes associated with ECM organization were prevalent, including Agt, Tnr, Myh11, Col5a3, Adamts14, Mmp21, and Adamtsl3. Additionally, we observed upregulation in genes involved in cell adhesion, such as Cd80, Tespa1, Mmrn1, Muc4, Gli3, Ptpru, Ripor2, Cd44, Vcam1, and Bmp6. And genes of components of the basement membrane, specifically Lama1 and Col5a3, were also upregulated.

DoRothEA analysis, which describes a gene regulatory network containing signed transcription factors (TF) and target gene interactions, provided further insight into the molecular mechanism at the regulon level based on all the DEGs, and found the target genes of Myc, Hif1a, Ar, Klf5, and Screbf1 are most downregulated, while Arid3a, Thap11, and Bhlhe40 were most upregulated. The gene expression data from EMCN^-/-^ glomeruli endothelium is consistent with the reduction of endothelial cellular homeostasis and barrier function, increase in extracellular remodeling/thickening and cell adhesion, and genotype that is involved in maintaining vascular homeostasis in the context of normal GFB function. Due to this, we further validated the mRNA expression eNOS and angiopoietin–Tie pathway through qPCR (Supplemental Figure 4 AB). We found EMCN^-/-^ kidneys show significant decrease in the transcriptional levels of eNOS (0.4821 ± 0.265 vs. 1.368 ± 0.248, *p* < 0.005, Student’s t-test), Tie2 (0.756 ± 0.059 vs. 1.023 ± 0.086, *p* < 0.005, Student’s t-test).

### Loss of EMCN in the endothelium leads to gene expression changes in glomerular podocytes that are involved in maintaining the structure and function of the GFB

Kidneys of EMCN^-/-^ mice exhibited disrupted GFB function. Podocytes, which play a crucial role in maintaining the GFB, showed effaced and fused foot processes in EMCN^-/-^ mice. Previous studies have reported that podocyte injury is associated with increased albuminuria under pathological conditions, such as in hyperglycemia^29,30^. To understand how the loss of EMCN also impacts podocytes, we isolated and performed RNA-seq on the podocyte population, as shown in Figures 7A and B. A total of 216 DEGs were found in EMCN^-/-^ glomeruli podocytes compared to EMCN^+/+^ controls, as shown in the volcano plot (Figure 8A). Of these, 87 genes were downregulated, while 129 genes showed upregulated expression. A heatmap of top-regulated DEGs in podocyte is shown in Figure 8B. DoRothEA analysis of all the DEGs in podocytes (Figure 8C) found the target genes of Klf6, Tead1, Ebf1, Myc and Screbf1 to be the most downregulated, and target genes of Hnf4g, Zfp639, Hnf1a, Stat4 and Thap11 are most upregulated.

**Figure 8.**
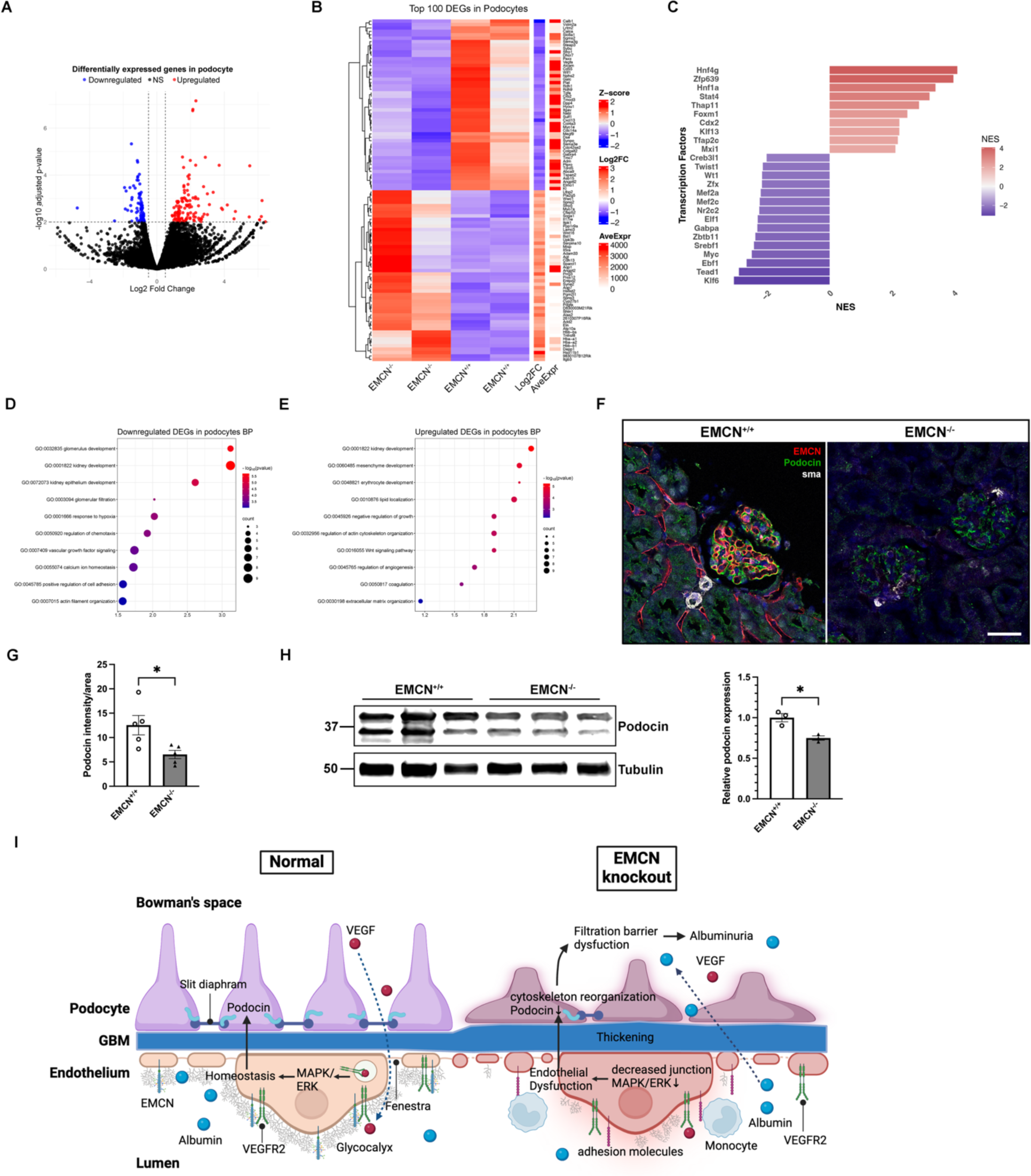
Kidney podocyte RNA-seq with differentially expressed genes analysis and schematic illustration of changes in EMCN knockout mice. **(A)** Volcano map of differentially expressed genes in podocytes in EMCN^-/-^ kidney endothelium compared to EMCN^+/+^. *p* < 0.01 with log2FC > 0.5 is considered as significantly expressed genes. Blue and red dots indicated downregulated and upregulated genes in EMCN^-/-^ kidney podocytes respectively. **(B)** Heatmap of top 100 DEGs with basemean>50 in EMCN^-/-^ kidney podocytes compared to EMCN^+/+^. **(C)** DoRothEA regulons analysis of transcription factor from DEGs in EMCN^-/-^ kidney podocytes compared to EMCN^+/+^. **(D)** Gene ontology biological process analysis of downregulated genes and **(E)** upregulated genes in EMCN^-/-^ kidney podocytes compared to EMCN^+/+^. **(F)** Immunohistochemical staining was performed to visualize the expression of EMCN, podocin, and smooth muscle actin. Nuclei were stained with DAPI. n = 3, Scale bar = 50 μm. **(G)** Podocin intensity per glomeruli area was quantified using ImageJ software. n = 5, * *p* < 0.05, Student’s t-test **(H)** Kidney protein samples were collected from EMCN^+/+^ and EMCN^-/-^ mice and quantified by BCA protein assay kit. Protein (40 µg) samples from three animals were loaded on an SDS-PAGE gel. Levels of podocin and tubulin were examined by western blot analysis. Band intensities of podocin from western blot were quantified by ImageJ software and normalized to tubulin. n = 3, * *p* < 0.05, Student’s t-test. **(I)** Schematic illustrating the proposed mechanism of kidney defects and dysfunction in EMCN^-/-^ mice, indicating that the loss of EMCN leads to endothelial dysfunction and filtration barrier impairment which results in the albuminuria observed in EMCN^-/-^ mice.

From the heatmap of top DEGs in podocytes (Figure 8B) and the GO analysis of biological process of the downregulated genes (Figure 8D), we noted a downregulation of genes involved in kidney and glomerulus development, and glomerular filtration, like Calb1, Sfrp1, Ptpro, Col4a3, Sulf1, Vegfa, Myo1e, Enpep, Robo2. We observed downregulation of genes associated with VEGF signaling and hypoxia, such as Sema3e, Lrtm2, Ptpro, Sema3g, Alcam, Vegfa, Robo2, Pak1, Slc8a1, Dpp4, Hyou1, and Plat. Additionally, genes associated with actin filament organization such as Sema3e, Nphs2, Gdpd2, and Pak1 were found downregulated. Validation of Nphs2 (podocin) downregulation at mRNA level was shown in Supplemental Figure 4C. GO analysis of biological processes (Figure 8E) of upregulated DEGs in podocytes revealed an increase in genes associated with mesenchyme development, including Ranbp3l, Heyl, Wnt7b, Sema4d, Frzb, Mdk, Wnt16, Wnt11, and Grem1. Additionally, there was upregulation in genes involved in actin cytoskeleton organization, such as Ppp1r9a, Capg, Itgb3, Mdk, Bst1, and Wnt11. Nhps2, also referred to as podocin, a component of the podocyte filtration slits that serves a critical function in maintaining the GFB ^31,32^, was downregulated in podocytes. We validated the downregulation of podocin at mRNA level through qPCR (1.009 ± 0.085 vs. 0.7366 ± 0.076, p < 0.05, Student’s t-test) (Supplemental Figure 4C). Examination by immunohistochemistry revealed reduced podocin staining intensity in the glomeruli of EMCN^-/-^ mice compared to EMCN^+/+^ (Figure 8F), which was confirmed by quantification of podocin staining intensity per glomeruli area (6.528 ± 0.844 pixels vs 12.54 ± 1.982 pixels, *p* < 0.05, Student’s t-test) (Figure 8G). These results were further supported by quantification of podocin by western blot affirming a significant decrease in protein levels of podocin in kidneys from EMCN^-/-^ mice compared to that from EMCN^+/+^ mice (Figure 8H) (0.750 ± 0.0268 vs 1.000 ± 0.0505, *p* < 0.05, Student’s t-test).

## DISCUSSION

The glycocalyx has been shown to play a significant role in regulating vascular homeostasis, and various pathophysiological processes^4,33^. Due to the widespread expression of EMCN^17^ and our previous in vitro and in vivo findings that EMCN prevents leukocyte-endothelial interactions under non-inflammatory conditions, its critical role in modulating VEGFR2 signaling in ECs, and during developmental angiogenesis^15,20^, we suspected that global EMCN KO mice might be embryonic lethal. Thus, we were surprised that EMCN^-/-^ mice were viable, born at the expected Mendelian frequency, and developed normally. While EMCN plays an important role in endothelial biology, its function during development appears to be compensated for in this genetic KO model^34^.

A comparison of expression levels of EMCN across a variety of tissues revealed relatively high levels of EMCN in kidneys. Thus, we explored the kidney phenotypes in adult EMCN^-/-^ mice. The kidneys are highly vascularized organs which receive up to 20% of cardiac output^35^. The kidney vasculature regulates the exchange of water and solutes between the blood and the interstitial fluid, which is essential for the maintenance of kidney function^36^. In kidneys, EMCN was primarily detected within the glomeruli, which are formed by unique bundles of capillaries lined by fenestrated endothelia and constitute a major component of the functional filtration unit of the kidney^35^. Consistent with our observation of EMCN expression in kidneys, a study of renal vascular development reported EMCN expression in embryonic glomeruli at E13 and increasing in level when examined at E16^37^.

We observed a significant increase in CD45^+^ cell infiltration, particularly monocytes, as well as an upregulation of VCAM-1 expression in the kidneys of EMCN^-/-^ mice. The finding of increased infiltrating inflammatory cells follows our previous observations that EMCN is involved in preventing leukocyte-endothelial interactions^14^, while increased VCAM-1 expression suggests a pro-inflammatory endothelial phenotype in the absence of EMCN. The role of the endothelial glycocalyx in preventing leukocyte-EC interactions has been demonstrated by many studies^38,39^ and reduced glycocalyx thickness is associated with increased leukocyte adhesion to ECs in the absence of fluid shear stress^38^. Degradation of the endothelial glycocalyx by local microinjection of heparinase significantly increases the number of adherent leukocytes^40^. The absence of podocalyxin in mice, another example of a highly sialoglycosylated glycocalyx glycoprotein, leads to the development of severe vasculitis and an increase in leukocyte adhesion^41^. In contrast, localized administration of key glycocalyx components such as heparan sulfate or heparin results in a significant reduction in leukocyte adhesion^3^. VCAM-1, which expression was elevated in the kidneys of EMCN null mice, is expressed by the endothelium and acts as a ligand for integrins found on leukocytes and platelets^2^, and facilitates the attachment of mononuclear leukocytes, such as monocytes and lymphocytes^42^. Increased VCAM-1 expression is detected in several vascular inflammatory diseases, including atherosclerosis^43^ and nephritis^44^, and therefore upregulation of VCAM-1 expression in the EMCN^-/-^ kidney is consistent with vascular inflammation of the glomeruli which could contribute to the functional defect of the GFB.

While no discernible differences were observed in kidney gross size or average glomerular count between EMCN^-/-^ and EMCN^+/+^ mice, a small but significant decrease in the average size of glomeruli in EMCN^-/-^ kidneys by H&E staining, along with increased collagen deposition, evidenced by trichrome staining, indicating basement membrane thickening and a fibrotic change of the kidney. Functional defects were indicated by the significant increase of albumin in the Bowman’s space and the urine of the EMCN^-/-^ mice compared to EMCN^+/+^ littermates. No significant difference in blood pressure changes was observed in the EMCN knockout, supporting the idea that the increased albuminuria is due to functional and/or structural changes in the GFB and is not caused by changes in blood pressure. Albuminuria is characteristic of various renal glomerular disorders^45^ and is commonly due to the disruption of the GFB^46^. The GFB, which includes the endothelium, glomerular basement membrane (GBM), and podocytes, controls the filtration of albumin and other circulating macromolecules^47^ and effectively reduces their levels in the urine. Examination of glomerular ultrastructure showed disorganized and thickened ECs, disorganized fenestrations, a thickened GBM, and effaced and fused podocytes. These observations are consistent with emerging evidence indicates that the endothelial glycocalyx is involved in regulating the filtration of albumin and other circulating macromolecules^47-49^.

To gain insight into the cell mechanism(s) involved in the structural and function change of GFB, we employed FACS on glomeruli endothelium and podocytes for RNA-seq. Results of these studies indicated reduced glycocalyx integrity and increased remodeling, as evidenced by lower levels of other glycocalyx components such as syndecan-4 and glypican-3 and elevated levels of matrix metalloproteinase (MMP) and ADAMTS proteases. Glypican-3 is a heparan sulfate-rich proteoglycan anchored to the cell surface known to bind growth factors, cytokines, and ECM components^2^. In the kidney, glypican-3 modulates bone morphogenic protein and fibroblast growth factor signaling during development^50^ and interacts with various signaling pathways, particularly Wnt, to influence cell proliferation and apoptosis in the adult^51^. Syndecan-4 is a transmembrane heparan sulfate-rich proteoglycan involved in cell adhesion, migration, and signal transduction^52^. It plays a key role in maintaining the structural and functional integrity of ECs through interaction with integrins and other ECM proteins, facilitating cell-matrix adhesion and mechanical stability of the glomerular endothelium^53,54^. ADAMTS proteases and MMPs play a critical role in ECM remodeling by proteolytically cleaving ECM components^55,56^.

Endothelial activation-induced glycocalyx degradation and enzymatic disruption of the glycocalyx^57^ or glycocalyx components^58^ have been shown to lead to albuminuria ^7^. For instance, conditional deletion of the enzyme core-1 β1,3 galactosyltransferase, which is critical for the synthesis of mucin-type core 1-derived O-glycans of all sialomucins by glomerular ECs and podocytes, results in impaired podocyte foot processes and spontaneous proteinuria^59^. Similarly, reduction of tight junction proteins and impaired cell adhesion dynamics indicated by RNA-seq suggests that the loss of EMCN disrupts the signaling pathways that maintain tight junctions, which would result in compromised barrier function, allowing for increased leakage of proteins and other molecules across the glomerular barrier^60^. We reported an essential role of EMCN in VEGFR2 endocytosis and downstream VEGF signaling^61^. The endothelial RNA-seq data, indicate the downregulation of MAPK/ERK signaling cascade and protein phosphorylation, which are not only downstream of the VEGF signaling pathway^62^ but also critical for cellular responses to stress and injury^63^. Additionally, the RNA-seq analysis of EMCN^-/-^ ECs shows increased expression of laminin-1 and collagens, which aligns with the thickened basement membrane observed in our TEM images. Besides, we found a decrease in eNOS, Tie2 expression in the EMCN^-/-^ kidneys, which are key markers of endothelial dysfunction^64-66^. Due to the limited number of cells sorted from fresh kidneys by FACS, we could only validate mRNA expression in bulk RNA extract from the renal cortex, whereas eNOS and Tie are highly specific to the endothelium^67,68^. VEGF induces eNOS phosphorylation through the PI3K-Akt signaling pathway^69^, which leads to prompt generation of NO^70^. Changes in the angiopoietin/Tie-2 axis are also closely associated with glomeruli endothelial dysfunction and albuminuria^71-73^. The DoRothEA analysis result further support the endothelium changes, as Myc, Hif1a and Klf5 consistently indicated the downregulation of endothelial cell proliferation, growth, and metabolism especially during hypoxia and inflammation^74-76^.

Podocytes play an essential role in preserving the selectivity of the GFB^45,77,78^. Although EMCN is predominantly expressed in endothelial cells, the crosstalk between endothelial cells and podocytes is well-documented, which means that the actions of one cell type can significantly impact the function of the other^29,79^. Selective damage of podocytes through saponin injection in Bowman’s space leads to albuminuria in vivo^80^. Podocyte injury, presented as foot process effacement and podocyte loss, are common to glomerulopathies with proteinuria^81^. The RNA-seq results of podocytes revealed downregulation of genes associated with kidney and glomerular filtration function such as Ptpro (also known as GLEPP1, glomerular epithelial protein 1), which is a receptor-type protein tyrosine phosphatase that plays a crucial role in podocyte function. Ptpro-deficient mice exhibit significant alterations in podocyte structure, including broader and flattened foot processes, and are associated with a decrease in glomerular filtration rate^82^. Wt1 (Wilms’ tumor 1), a master regulator of gene expression in podocytes, controls the production of several podocyte-specific proteins including podocalyxin, a major membrane protein essential for podocyte structure and function^83^. The podocyte RNA-seq results also revealed downregulation of the VEGF signaling pathway and hypoxia response. Produced by podocytes, VEGF acts both in an autocrine manner to promote podocyte survival and as a paracrine factor to maintain the fenestrated phenotype of adjacent glomerular ECs, which is essential for proper filtration^84,85,86^. This role for VEGFA is a possible explanation for our findings in EMCN^-/-^ kidneys where impaired VEGFR2 signaling would be expected due to the absence of EMCN^15,20^. Consistent with this concept, long-term follow-up of patients receiving intravitreal injections of VEGF inhibitors, such as aflibercept, ranibizumab, and bevacizumab exhibit evidence of systemic absorption^87,88^ and have presented with manifestations of nephrotoxicity, including hypertension and albuminuria^66, 89^. Furthermore, proteinuria is commonly observed as a detrimental vascular effect in individuals with advanced-stage solid tumors who undergo anti-angiogenic treatment specifically targeting VEGF and/or its receptors^90-93^.

Notably, genes involved in actin cytoskeleton organization and stabilization were altered with EMCN KO in podocytes. Capg (capping protein, gelsolin like) and Itgb3 (integrin beta-3) are involved in actin filament dynamics and cell adhesion, respectively, playing critical roles in podocyte stability and function^94,95^. Nphs2 encodes podocin, a crucial component of the slit diaphragm that localizes to the foot process membrane^32^ and constitutes the filtration diaphragm in podocytes^96^. Podocin colocalizes to tight junctions between foot processes and serves as scaffolding tethering tight junction proteins to the cytoskeleton^97^. Its reduction is a biomarker for early podocyte injury and dysfunction^98,99^. Podocin mutations are found both in hereditary and sporadic glomerulosclerosis, which are associated with proteinuria^100-102^. Proteolytic cleavage of podocin by Matriptase, a podocyte protein activated in patients and mice with chronic kidney diseases, exacerbates podocyte injury^103^. Proteinuria is most often thought to result from alterations in podocyte foot processes. However, increasing evidence shows glomeruli endothelial abnormalities^104,105^, with or without podocyte changes^106,107^, are also closely associated with increasing urine albumin excretion, suggesting that EC-podocyte crosstalk is critical to GFB function^30,108,109^. An important avenue for future research would be to examine EMCN’s function in models of acute or chronic kidney injury. Such studies could reveal EMCN’s role during renal stress, complement our current findings, and provide a more comprehensive understanding of the significance of glycocalyx in disease.

A schematic illustration summarizes the potential mechanism underlying the pathological changes observed in kidneys lacking EMCN is shown in Figure 8I. Concisely, under normal conditions, endothelial-podocyte crosstalk maintains homeostasis and GFB function. The paracrine VEGF from podocytes passes through the basement membrane to act on ECs. Homeostatic endothelium modulates the permeability of albumin passing through the GFB. Genetic deletion of EMCN, which closely regulates VEGFR2 endocytosis and VEGF signaling as a component of the glycocalyx, disrupts glycocalyx integrity and glomeruli endothelial tight junctions potentially through MAPK/ERK pathways, and increases infiltration of CD45^+^ cells, especially monocytes, was observed in the in kidney. Disruption of glomerular endothelial homeostasis combined with inflammation further impacts podocytes, leading to repression of genes maintaining podocyte function, like VEGF signaling and Wt1, and disruption of actin cytoskeleton organization, which maintains foot process and slit diaphragm function and contributes to dysfunctional GFB and albuminuria in EMCN^-/-^ mice. Taken together, these observations indicate that the absence of EMCN leads to a functional defect in the GFB and results in increased serum albumin in the Bowman’s space and urine.

Our phenotypic analysis of the first global EMCN deletion establishes EMCN’s critical role in the maintenance of the GFB. This suggests a crucial role for the endothelial glycocalyx in maintaining the glomerular structure and function, especially in EC-podocyte crosstalk by modulating VEGF signaling and suppressing endothelial inflammation.

## Acknowledgments

The authors thank Philip Seifert at The Schepens Eye Research Institute of Mass. Eye and Ear Morphology Core and Diane E. Capen at the Massachusetts General Hospital Center for Systems Biology and Program in Membrane Biology/Division of Nephrology for TEM. We appreciated the Flow Cytometry Core Facility at Schepens Eye Research Institute of Mass. Eye and Ear and the HSCI-CRM Flow Cytometry Core Facility at Mass General Hospital. We appreciate Dr. Dennis Brown from Mass General Hospital Membrane Biology, Division of Nephrology, and the Center for Systems Biology for providing suggestions and guidance on urinalysis.

## NOVELTY AND SIGNIFICANCE

### What is known?

- EMCN, an endothelial-specific component of the vascular glycocalyx, also distinguishes hematopoietic stem cells from lineage-committed hematopoietic progenitor cells.
- EMCN plays a role in preventing leukocyte-endothelium adhesion and regulating VEGF-induced VEGFR2 endocytosis and signaling in vitro.

### What new information does this article contribute?

- Mice with global knockout of EMCN are viable.
- Lack of EMCN leads to increased CD45^+^ cell infiltration, particularly ly6G^low^ly6C^high^ monocytes/macrophage lineage cells in kidney glomeruli.
- Loss of EMCN results in disrupted GFB function, with ultrastructural defects of the glomeruli, including disorganized endothelium, thickened GBM, as well as effaced and fused podocyte foot processes.
- The absence of EMCN likely disrupts the glomerular filtration barrier by compromising endothelial glycocalyx and tight junction integrity, contributing to increased immune cell infiltration, endothelial and podocyte dysfunction
- Leukocyte infiltration is further facilitated by the lack of endothelial cell surface EMCN, which has been shown to maintain a non-inflammatory cell surface

The endothelial glycocalyx, which lines the luminal surface of the vascular endothelium, is essential to maintaining the integrity of the endothelial barrier. The function of EMCN, a component of the endothelial glycocalyx selectively expressed in the capillaries and venous endothelium, has not been fully explored. In this study, we generated global EMCN knockout mice. Motivated by the high expression of EMCN in the kidney, we investigated the renal kidney phenotypes in EMCN^-/-^ mice. We observed increased infiltration of inflammatory cells, particularly monocytes in EMCN^-/-^ kidneys. Additionally, EMCN^-/-^ mice exhibited albuminuria accompanied by a thickened basement membrane, disrupted and disorganized fenestrated endothelium, as well as podocyte effacement. The absence of EMCN was found to disrupt endothelial homeostasis, and together with the infiltration of inflammatory cells, ultimately leads to dysfunction of the glomerular filtration barrier and the subsequent albuminuria. These findings shed light on the critical role of the glycocalyx molecule EMCN in maintaining kidney vascular permeability and the integrity of the glomerular filtration barrier.

---

**Supplemental Figure 1.**
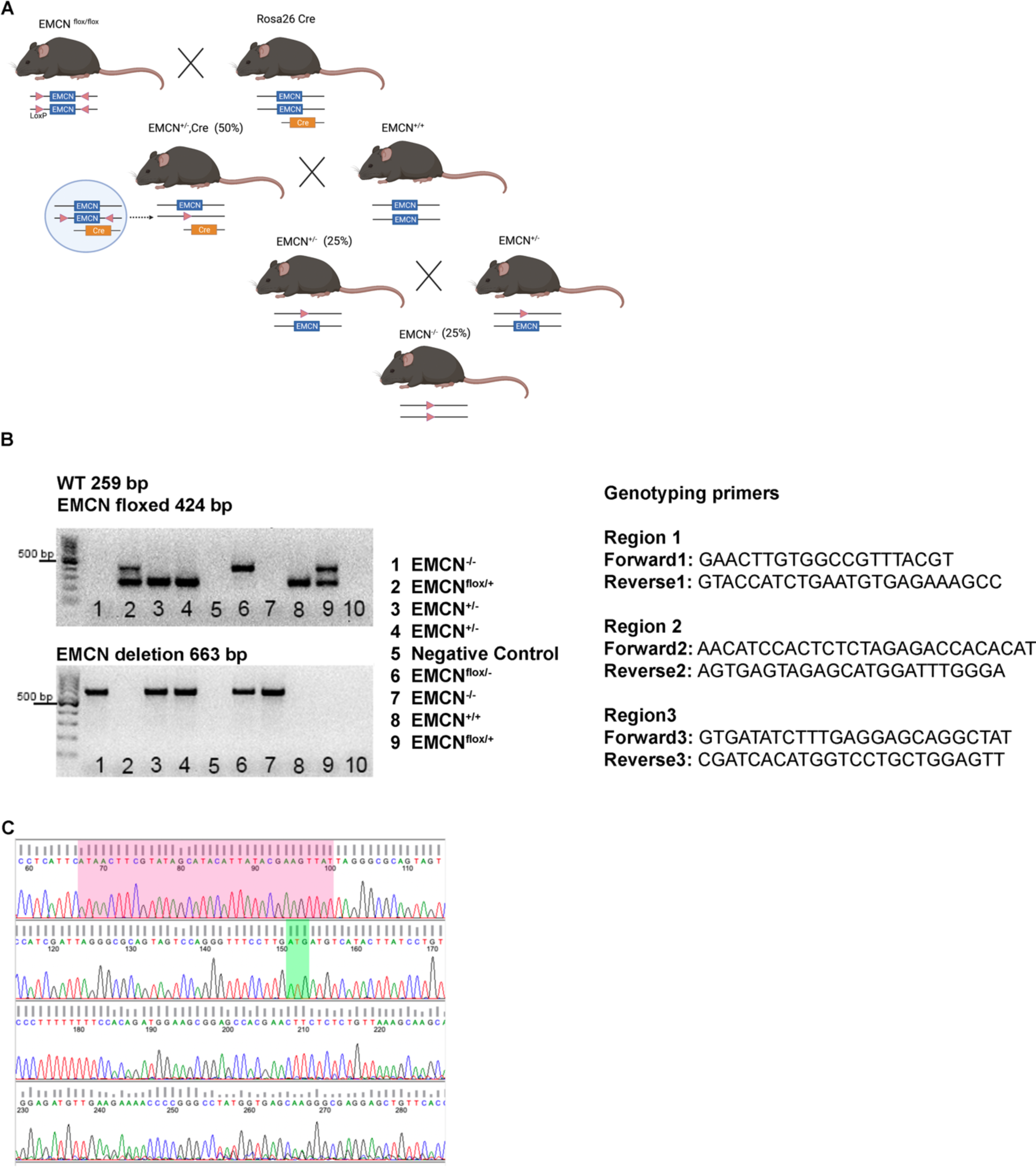
Generation and confirmation of EMCN^-/-^ mice. **(A)** The image illustrates the step-by-step strategy for the generation of global ^-/-^ mice. **(B)** A representative image displaying the genotyping result obtained by PCR amplification of region 2 using genomic DNA from EMCN wild-type, EMCN floxed, and EMCN deletion alleles. The primers utilized for PCR amplification are provided in the list. **(C)** The Sanger sequencing confirms the deletion of the EMCN sequence from an EMCN deletion allele. The Loxp site sequence is indicated in pink, while the start codon for EGFP is labeled in green.

**Supplemental Figure 2.**
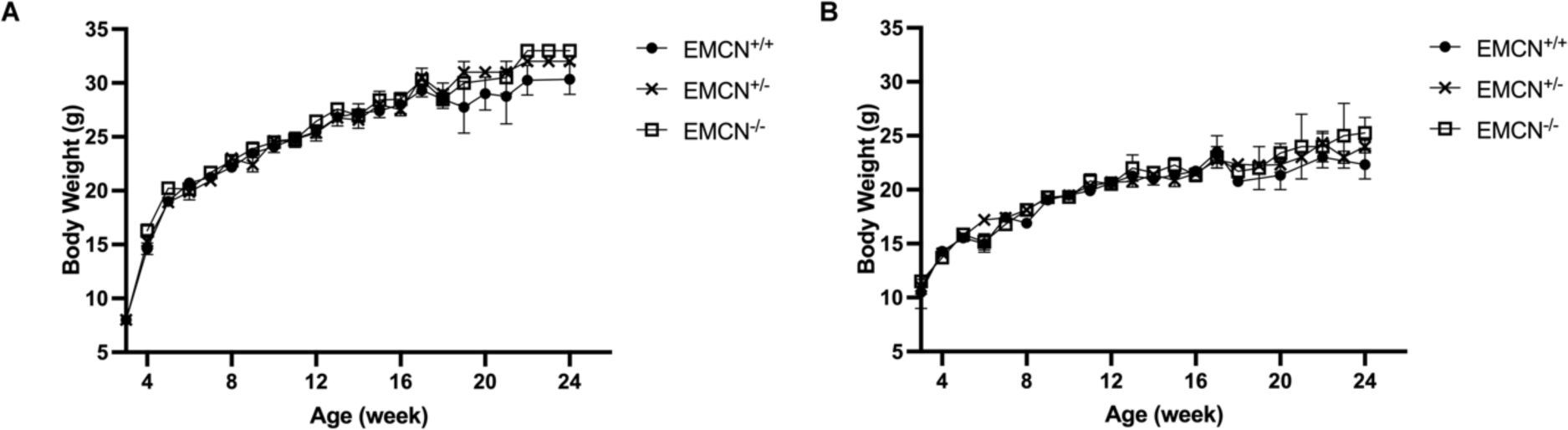
Body weight of EMCN^+/+^, EMCN^+/-^, and EMCN^-/-^ mice up to 6-months-old. Body weights of EMCN^+/+^, EMCN^+/-^, and EMCN^-/-^ **(A)** male and **(B)** female mice were tracked from week four to week 24. n > 5 for all time points and genotypes. *p* > 0.05, one-way ANOVA.

**Supplemental Figure 3.**
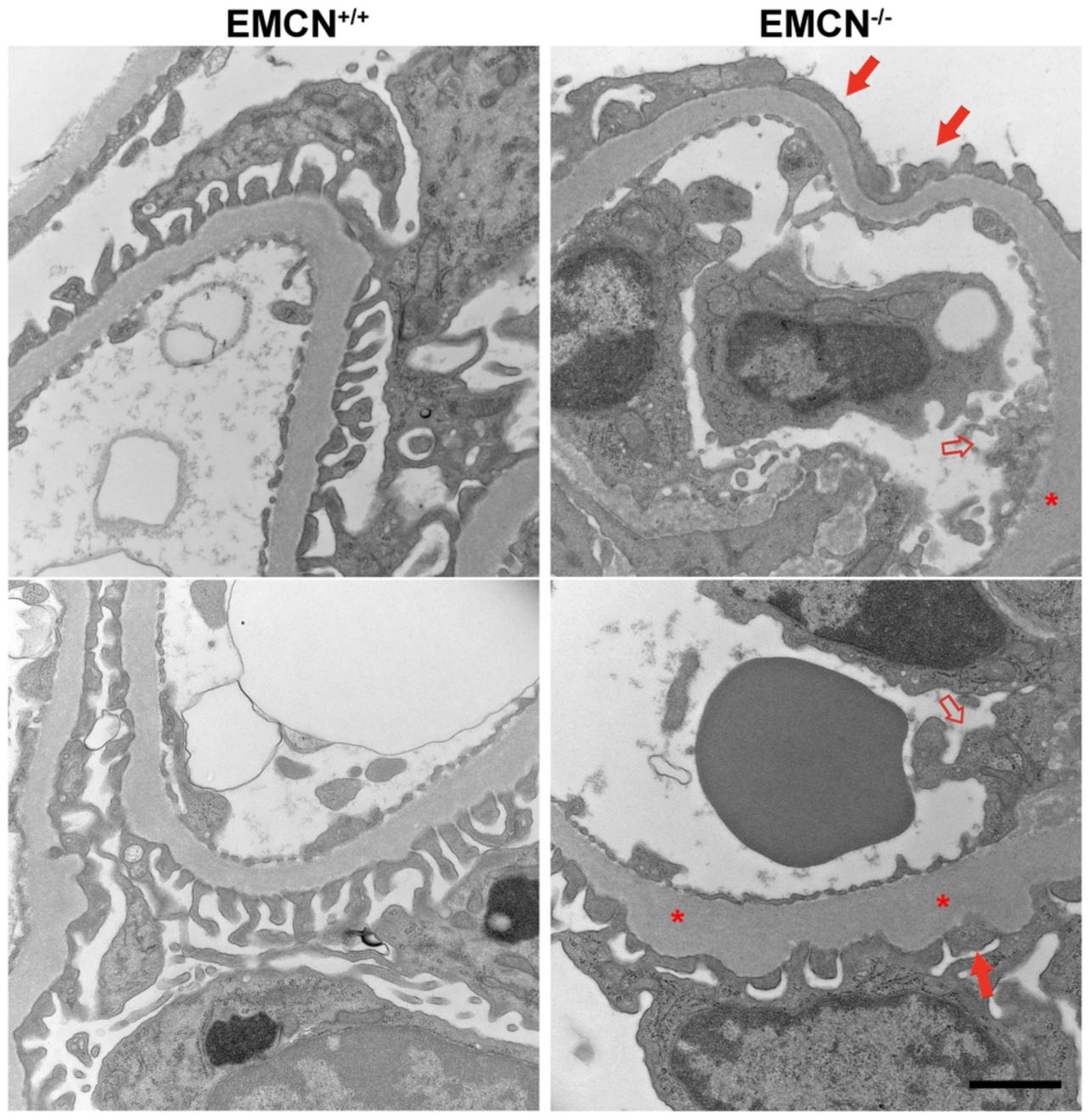
Glomerular ultrastructure in kidneys from aged EMCN^-/-^ mice. Representative TEM images of the GFB of 21-24-month-old littermates of EMCN^+/+^ (left panels) and EMCN^-/-^ (right panels). Red arrows indicate effaced and fused podocytes, red hollow arrows indicate disorganized endothelial fenestrations, and red asterisks indicate basement membrane thickening in EMCN^-/-^ mice. n = 2. Scale bar = 1 μm.

**Supplemental Figure 4.**
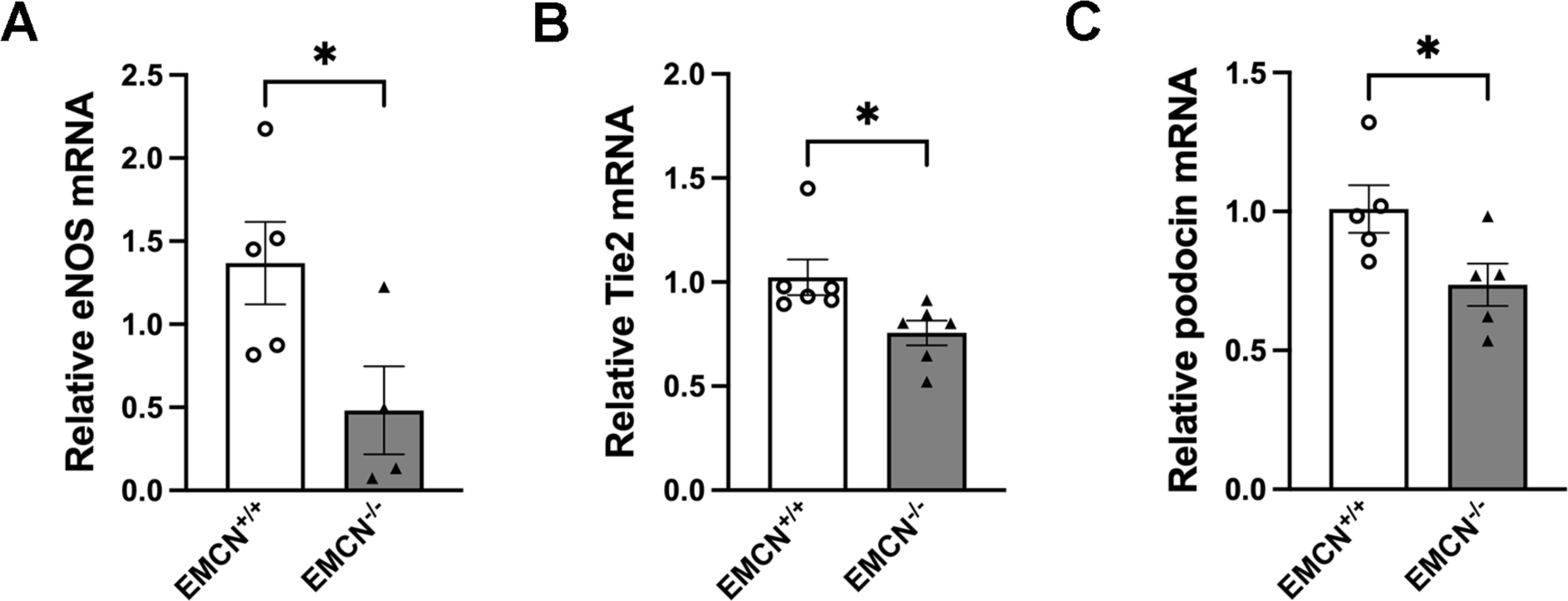
Validation of mRNA changes in eNOS, Tie2, and podocin. Total RNA was extracted from kidneys of EMCN^+/+^ and EMCN^-/-^ mice and analyzed for mRNA expression of **(A)** eNOS, **(B)** Tie2, and **(C)** podocin. * *p* < 0.05, Student’s t-test.

